# Alzheimer’s disease-associated *TM2D* genes regulate Notch signaling and neuronal function in *Drosophila*

**DOI:** 10.1101/2021.04.20.440660

**Authors:** Jose L. Salazar, Sheng-An Yang, Yong Qi Lin, David Li-Kroeger, Paul C. Marcogliese, Samantha L. Deal, G. Gregory Neely, Shinya Yamamoto

## Abstract

TM2 domain containing (TM2D) proteins are conserved in metazoans and encoded by three separate genes in each species. Rare variants in *TM2D3* are associated with Alzheimer’s disease (AD) and its fly ortholog *almondex* is required for embryonic Notch signaling. However, the functions of this gene family remain elusive. We knocked-out all three *TM2D* genes (*almondex, CG11103/amaretto*, *CG10795/biscotti*) in *Drosophila* and found that they share the same maternal-effect neurogenic defect. Triple null animals are not phenotypically worse than single nulls, suggesting these genes function together. Overexpression of the most conserved region of the TM2D proteins acts as a potent inhibitor of Notch signaling at the γ-secretase cleavage step. Lastly, Almondex is detected in the brain and its loss causes shortened lifespan accompanied by progressive electrophysiological defects. The functional links between all three *TM2D* genes are likely to be evolutionarily conserved, suggesting that this entire gene family may be involved in AD.

## Introduction

Alzheimer’s disease (AD) is the most common neurodegenerative disease affecting the aging population and accounts for the large majority of age-related cases of dementia (Long and Holtzman, 2019; Sala Frigerio and De Strooper, 2016). AD is pathologically characterized by histological signs of neurodegeneration that are accompanied by formation of extracellular plaques and intra-neuronal tangles. Numerous studies have identified genetic factors that contribute to AD risk and pathogenesis (Bellenguez et al., 2020). Rare hereditary forms of AD are caused by dominant pathogenic variants in *APP (Amyloid Precursor Protein)*, *PSEN1* (*Presenilin 1*) or *PSEN2 (Presenilin 2)*. These three genes have been extensively studied using variety of experimental systems, and the resultant knowledge has led to greater understanding of how they contribute to the formation of extracellular plaques found in both familial and sporadic AD brains (Karran et al., 2011). *PSEN1* and *PSEN2* are paralogous genes that encode the catalytic subunit of the γ-secretase, a membrane-bound intramembrane protease complex (Zhang et al., 2013). γ-secretase substrates include many type-I transmembrane proteins including APP as well as Notch receptors that play various roles in development and physiology (Artavanis-Tsakonas et al., 1995; Kopan and Ilagan, 2009). Processing of APP by γ-secretase generates small peptide fragments of varying length, collectively termed amyloid beta (Aβ). The plaques characteristic of AD are largely composed of Aβ peptides, likely seeded by the most oligomeric and neurotoxic species of Aβ that is 42 amino acids long (Aβ42). Since deposition of amyloid plaques can be found in pre-symptomatic stages of the disease (Sperling et al., 2011), many consider the production of toxic Aβ peptides to play a critical role during the very early phase of AD pathogenesis. Interestingly, individuals with one duplicated copy of wild-type APP including Down syndrome patients (trisomy 21, *APP* is on chromosome 21) have significantly increased risk and earlier age-of-onset of AD (Lott and Head, 2019), indicating that simply increasing APP and its cleavage products is sufficient to increase AD risk. Increases in Aβ production can also trigger the formation of intra-neuronal tangles composed of hyper-phosphorylated Tau, which can directly impact neuronal function and mediate degeneration (Ballatore et al., 2007).

While studies of genes that cause familial AD have been critical in providing a framework to study pathogenic mechanisms of AD, pathogenic variants in *APP* and *PSEN1/2* are responsible for only a small fraction of AD cases (Cacace et al., 2016). Familial AD can be distinguished from more common forms of AD because most patients with *APP* or *PSEN1/2* variants develop AD before the age of 65 [early-onset AD (EOAD)]. The majority (>95%) of AD cases are late-onset (LOAD, develops after 65 years of age) and of sporadic or idiopathic nature (Goldman et al., 2011). In these patients, it is thought that multiple genetic and environmental factors collaborate to cause damage to the nervous system that converges on a pathway that is affected by APP and PSEN1/2. To reveal common genetic factors with relatively small effect sizes, multiple genome-wide association studies (GWAS) have been performed and have identified over 40 loci throughout the genome that confer increase risk to developing AD (Bellenguez et al., 2020). The most notable risk-factors are variant alleles in *APOE* (Strittmatter et al., 1993). Although the precise molecular mechanism by which different alleles of *APOE* increase or decrease the risk of AD has been extensively debated, a number of studies have proposed that this gene is involved in the clearance of toxic Aβ peptides (Serrano-Pozo et al., 2021). A recent meta-analysis has also identified *ADAM10* (encoding a β-secretase enzyme that cleaves APP and Notch) as an AD associated locus (Kunkle et al., 2019), suggesting that genes involved in familial EOAD and sporadic LOAD may converge on the same molecular pathway. Functional studies of these and other newly identified risk factors for AD are critical to fully understand the etiology of this complicated disease that lack effective treatments or preventions.

We have previously reported that a rare missense variant (rs139709573, NP_510883.2:p.P155L) in *TM2D3* (*TM2 domain containing 3*) is significantly (OR=7.45, pMETA =6.6×10^-9^) associated with increased risk of developing LOAD through an exome-wide association analysis in collaboration with the CHARGE (Cohorts for Heart and Aging Research in Genomic Epidemiology) consortium (Jakobsdottir et al., 2016). This variant was also associated with earlier age-at-onset that corresponds to up to 10 years of difference with a hazard ratio of 5.3 (95% confidence interval 2.7-10.5) after adjusting for the *ε4* allele of *APOE*. Although the function of this gene in vertebrates was unknown and this missense variant was not predicted to be pathogenic based on multiple variant pathogenicity prediction algorithms including SIFT (Sim et al., 2012), PolyPhen (Adzhubei et al., 2010) and CADD (Kircher et al., 2014), we experimentally demonstrated that p.P155L has deleterious consequences on TM2D3 function based on an assay we established using *Drosophila* embryos (Jakobsdottir et al., 2016). The *Drosophila* ortholog of *TM2D3*, *almondex* (*amx*), was initially identified based on an X-linked female sterile mutant allele (*amx^1^*) generated through random mutagenesis (Shannon, 1972). Although homozygous or hemizygous (over a deficiency) *amx^1^* mutant females and hemizygous (over Y chromosome) males are viable with no morphological phenotypes, all embryos laid by *amx^1^* hemi/homozygous mothers exhibit severe developmental abnormalities including expansion of the nervous system at the expense of the epidermis (Lehmann et al., 1983; Shannon, 1973). This ‘neurogenic’ phenotype results when Notch signaling mediated lateral inhibition is disrupted during cell-fate decisions in the developing ectoderm (Lewis, 1996; Salazar and Yamamoto, 2018). By taking advantage of this scorable phenotype, we showed that the maternal-effect neurogenic phenotype of *amx^1^* hemizygous females can be significantly suppressed by introducing the reference human TM2D3 expressed under the regulatory elements of fly *amx*, but TM2D3^p.P155L^ expressed in the same manner fails to do so (Jakobsdottir et al., 2016). This showed that the function of *TM2D3* is evolutionarily conserved between flies and humans, and the molecular function of *TM2D3* that is relevant to LOAD may also be related to Notch signaling. More recently, another rare missense variant (p.P69L) in this gene has been reported in a proband that fit the diagnostic criteria of EOAD or frontotemporal dementia (Cochran et al., 2019), indicating that other *TM2D3* variants may be involved in dementia beyond LOAD.

TM2D3 is one of three highly conserved TM2 domain containing (TM2D) proteins encoded in the human genome. The two other TM2 domain-containing proteins, TM2D1 and TM2D2 share a similar protein domain structure with TM2D3, and each protein is encoded by a highly conserved orthologous gene in *Drosophila* that have not been functionally characterized (*CG10795* and *CG11103,* respectively) (**Supplemental Figure 1**). All TM2D proteins have a predicted N-terminal signal sequence and two transmembrane domains that are connected through a short intracellular loop. Within this loop, there is an evolutionarily conserved DRF (aspartate-arginine-phenylalanine) motif, a sequence found in some G-protein coupled receptors that mediates their conformational change upon ligand binding (Koenen et al., 2017). The extracellular region between the signal sequence and first transmembrane domain is divergent in different species as well as among the three TM2D containing proteins. In contrast, the sequences of the two transmembrane domains as well as the intracellular loop is highly conserved throughout evolution as well as between the three TM2 domain containing proteins (Kajkowski et al., 2001) (**Supplemental Figure 1**). The three proteins also have short C-terminal extracellular tails that are evolutionarily conserved but vary among the three proteins (e.g. TM2D1 has a slightly longer C’-tail than TM2D2 and TM2D3). The molecular functions of these conserved and non-conserved domains of TM2D proteins are unknown.

In this study, we generated clean null alleles of all three *Drosophila TM2D* genes using CRISPR/Cas9-mediated homology directed repair (HDR) and assessed their functions *in vivo*. Surprisingly, we found that *CG10795* (*TM2D1*) and *CG11103* (*TM2D2*) knockout flies are phenotypically indistinguishable from *amx* (*TM2D3*) null animals, displaying severe maternal-effect neurogenic phenotypes. We also generated double- and triple-knockout animals to determine whether these three genes have redundant functions in other Notch signaling dependent contexts during development. The triple-knockout of all *TM2D* genes did not exhibit any obvious morphological phenotypes but shared the same maternal-effect neurogenic phenotype similar to the single null mutants, suggesting these three genes function together. We also provide evidence that Amx functions on γ-secretase to modulate Notch signaling *in vivo*, and further uncover a previously unknown role of this gene in maintenance of neural function in adults.

## Results

### A clean null allele of *amx* fully recapitulates previously reported maternal-effect phenotypes

In a previous study, Michellod et al. reported the generation of a null allele of *amx* (*amx^m^*) by generating flies that are homozygous for a deletion that removes *amx* and several other genes [*Df(1)FF8*] and bringing back a genomic rescue construct for *Dsor1*, an essential gene that lies within this locus (Michellod et al., 2003). The authors reported that zygotic *amx^m^* [actual genotype: *Df(1)FF8; P(Dsor1^+S^)*] mutant flies have reduced eye size and nicked wing margin as adults, which are phenotypes that had not been reported in the classic *amx^1^* allele. Because the *amx^1^* allele is caused by a 5bp deletion that may still produce a product that encodes a portion of the N’-extracellular domain (*amx* is a single exon gene in *Drosophila*, hence will not be subjected to nonsense mediated decay), the authors concluded that *amx^1^* is a hypomorphic allele.

Since *Df(1)FF8* has not been molecularly characterized and it was uncertain whether all of the phenotypes attributed to *amx^m^* are due to loss of *amx* alone, we decided to generate a clean null allele of *amx* (hereafter referred to as *amx^Δ^*) using CRISPR/Cas9 technology (Bier et al., 2018; Knott and Doudna, 2018). We knocked-in a dominant wing color marker (*y^wing2+^)* to replace the coding sequence of *amx* using HDR (Li-Kroeger et al., 2018) (**Figure 1A**). Insertion of this cassette was screened by the presence of the visible marker (dark wings) in a *yellow* mutant background and the targeting event was molecularly confirmed via Sanger sequencing. We also confirmed the loss of *amx* transcript by RT-PCR (**Supplemental Figure 2**). Similar to *amx^1^*, homozygous *amx*^Δ^ females exhibit sterility and all embryos produced by these animals exhibit a neurogenic phenotype (**Figure 1C-E**). Both female fertility and neurogenic phenotypes of their progeny can be suppressed by introducing a 3.3 kb genomic rescue construct containing the *amx* locus **(Figure 1A)** (Jakobsdottir et al., 2016). Human TM2D3 expressed using the same regulatory elements has ∼50% activity of fly Amx **(Figure 1C)**, consistent with what we previously observed using *amx^1^* hemizygous females. We did not observe any morphological defects in the eye and wing of homozygous and hemizygous *amx*^Δ^ flies at all temperatures tested (between 18-29°C). We also looked at other tissues that are often affected when Notch signaling is defective including the notum and legs (Córdoba and Estella, 2020; Schweisguth, 2015) but we did not observe any morphological phenotypes in these tissues either. In summary, *amx^Δ^* is the first clean loss-of-function (LoF) allele of *amx* generated by CRISPR/Cas9, and phenotypically resembles the classic *amx^1^* allele rather than the *amx^m^* allele.

**Figure 1.**
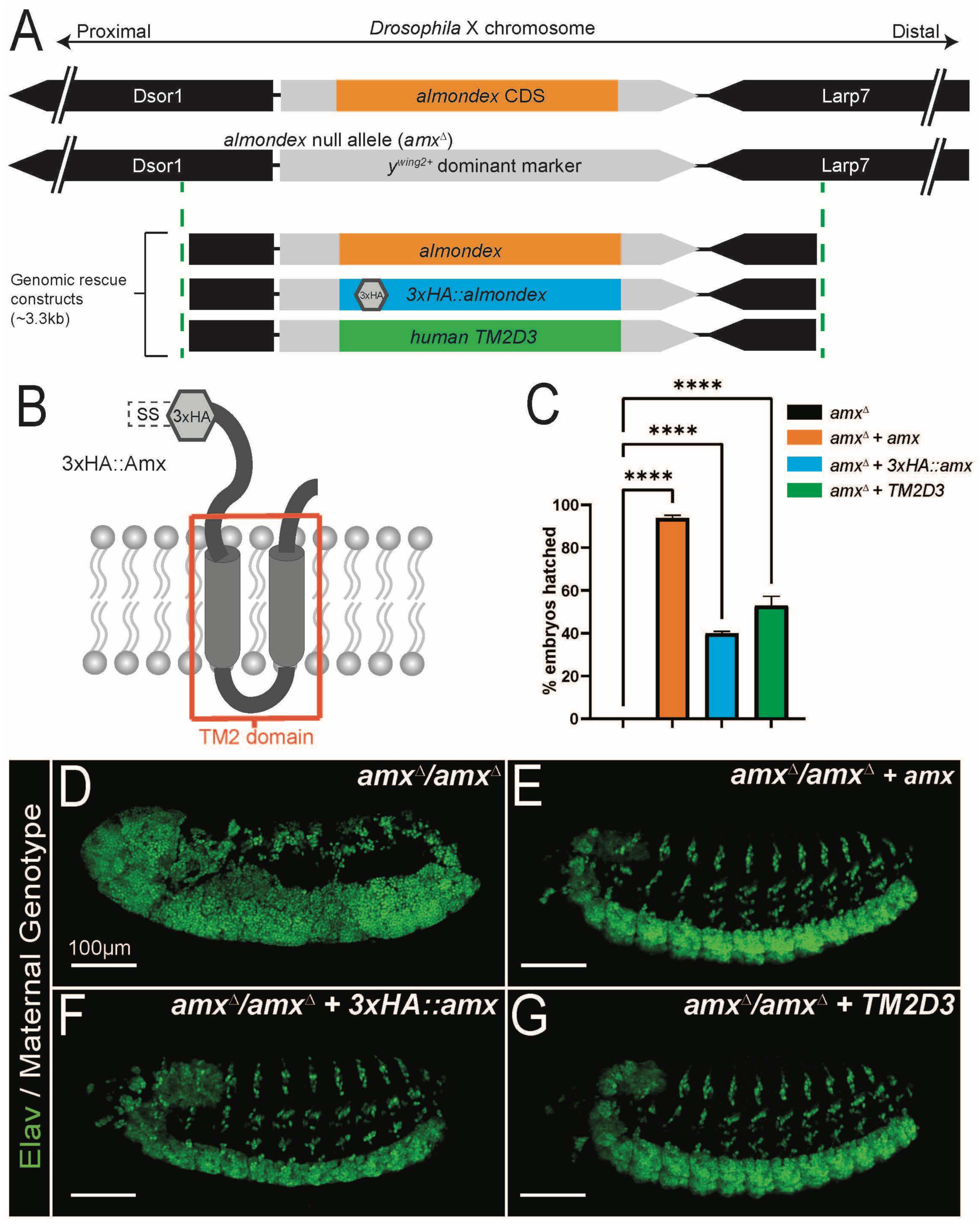
A clean null allele of *TM2D3* fly ortholog *almondex* (*amx^Δ^*) behaves like the classic *amx^1^* allele. (A) Schematic diagram of *almondex* (*amx*) locus, *amx^Δ^* allele and genomic rescue constructs used in this study. (B) Predicted 2D-structure of *Drosophila* Amx protein. SS = signal sequence for membrane localization. Transmembrane 2 (TM2) domain is boxed in red. Hexagon denotes where 3xHA epitope is located in 3xHA::Amx protein. (C) Egg hatching assay shows that genomic rescue constructs can suppress embryonic lethality (*amx^Δ^* n=857. *amx^Δ^* + *amx* n=139. *amx^Δ^* + *3xHA*::*amx* n=1673. *amx^Δ^* + *TM2D3* n=257). Error bars show SEM. One-way ANOVA followed by Dunnett test. **** = p-value≤0.0001. (D-G) Embryonic nervous tissue (neuronal nuclei, Elav, green) of developing embryos. Embryos from homozygous *amx^Δ^* females (D) exhibit a neurogenic phenotype. This phenotype can be suppressed by wild-type *amx* (E), *3xHA::amx* (F), or human *TM2D3* (G) genomic rescue constructs.

### Null alleles of other *TM2D* genes exhibit the maternal-effect neurogenic phenotype similar to *amx*

Using a similar strategy that we used to generate *amx^Δ^* (Li-Kroeger et al., 2018), we generated null alleles for *CG11103* (*TM2D2*, located on the X-chromosome) and *CG10795* (*TM2D1*, located on the 2^nd^ chromosome), two uncharacterized genes without official names (**Figure 2**). This time, we inserted a dominant body color marker (*y^body+^*) into the endogenous loci of *CG11103* and *CG10795* to knock-out these genes, and we phenotypically and molecularly characterized these alleles in a similar manner.

**Figure 2.**
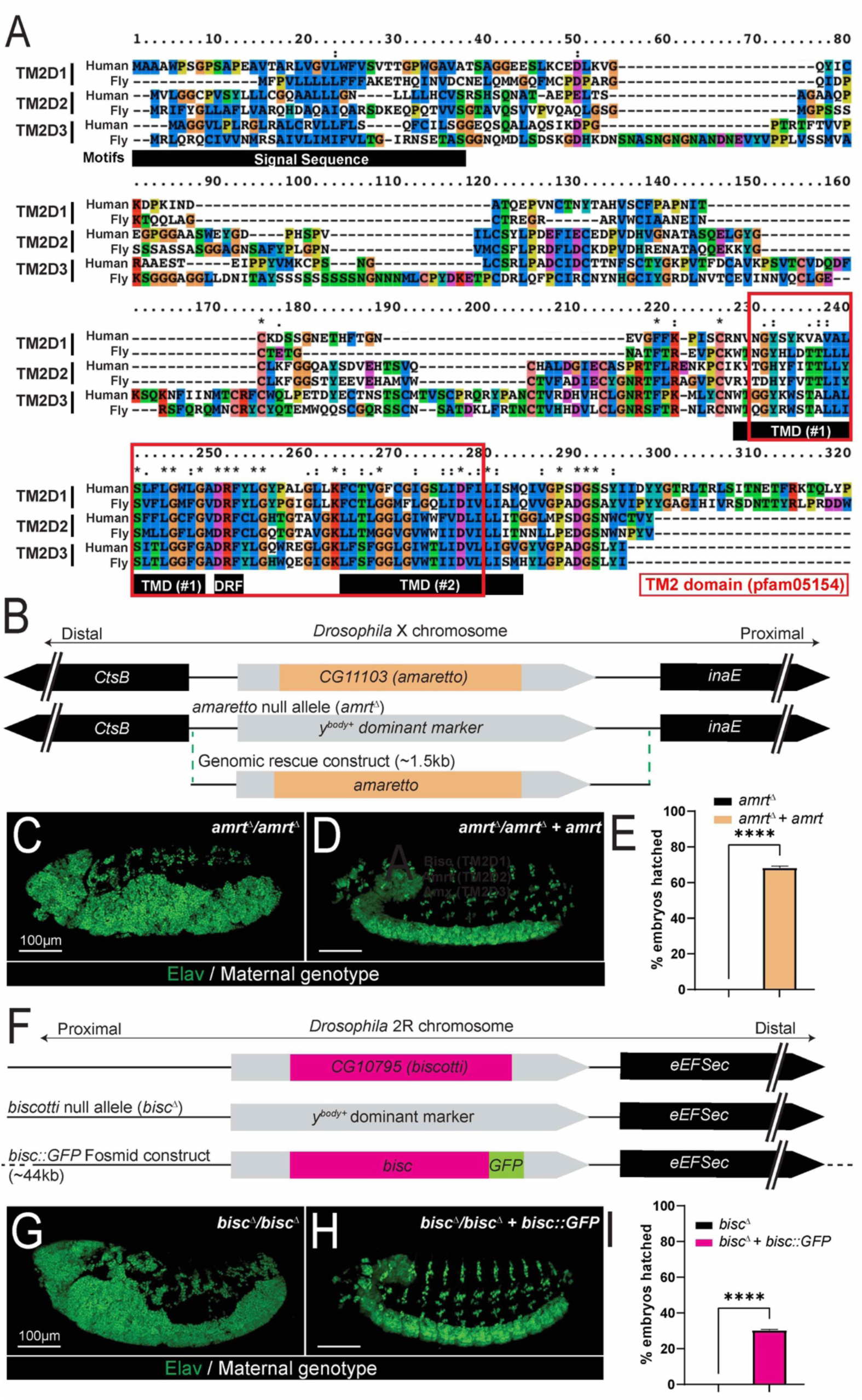
Null alleles of fly orthologs of *TM2D2* (*CG11103/amrt*) and *TM2D1* (*CG10795/bisc*) phenotypically mimics the loss of *TM2D3* (*amx*). (A) Protein alignment of human and *Drosophila* TM2D proteins. The TM2 domain (boxed in red) is composed of two transmembrane domains (TMD) and an intracellular DRF motif (denoted by black bars for TM2D3). (B) Schematic of *amaretto* (*amrt*) locus, *amrt* null allele (*amrt^Δ^*), and *amrt* genomic rescue plasmid construct generated for this study. (C) Embryos from homozygous *amrt^Δ^* females exhibit neurogenic phenotype, which can be suppressed by the *amrt* genomic rescue construct (D). (E) Egg hatching assay showing that *amrt* genomic rescue construct suppresses 68% of embryo lethality (*amrt^Δ^* n=954. *amrt^Δ^* + *amrt* n=1490). (F) Schematic of *biscotti* (*bisc*) locus, *bisc* null allele (*bisc^Δ^*), and *bisc::GFP* genomic rescue fosmid construct generated for this study. (G) Embryos from homozygous *bisc^Δ^* females exhibit neurogenic phenotype (n=379) which can be suppressed by *bisc::GFP* genomic rescue construct (H). (I) Egg hatching assay showing that *bisc::GFP* suppresses 30% of embryonic lethality (*bisc^Δ^* n=379*. bisc^Δ^ + bisc::GFP* n=436). t-test, **** = p-value<0.0001.

Both males and females that are hemizygous or homozygous null for *CG11103* are viable and these animals do not exhibit any morphological defects. Similar to *amx*, *CG11103* mutant females are fully sterile, and all embryos laid by these mothers exhibit neurogenic phenotypes (**Figure 2C**). Importantly, these phenotypes are rescued by a 1.5 kb genomic construct containing the *CG11103* locus (**Figure 2B, D, E**). Given the phenotypic and molecular similarities with *amx*, we gave *CG11103* the name *amaretto* (*amrt*), after the sweet Italian liqueur traditionally flavored with almonds. The knockout allele for this gene is referred to as *amrt^Δ^* hereafter.

Like *amx* and *amrt*, mutants that are homozygous null for *CG10795* appear morphologically normal, but all females are sterile and their embryos exhibit a neurogenic phenotype (**Figure 2G**). We were able to suppress these phenotypes using a ∼40kb fosmid transgene in which the CG10795 protein is C-terminaly tagged with multiple epitobes including GFP (Sarov et al., 2016). Given the phenotypic similarity to *amx* and *amrt* mutants, we named this gene *biscotti* (*bisc*), after the Italian biscuits traditionally made with almonds. The knockout allele for this gene is referred to as *bisc^Δ^* hereafter.

### Triple knockout of *TM2D* genes is phenotypically similar to single gene knockouts

Embryonic neurogenic defect is a rare phenotype that is almost exclusively associated with genes that affect the Notch signaling pathway (Lewis, 1996). Given the importance of Notch signaling in most stages of development preceding adulthood (Artavanis-Tsakonas et al., 1995), it was peculiar that all three *TM2D* gene mutants do not exhibit any obvious morphological defects related to Notch signaling while all three alleles exhibit a robust maternal-effect neurogenic phenotype. To test whether this may be due to redundancy between the three genes, we generated a fly strain that lacks all three *TM2D* genes (**Figure 3**).

**Figure 3.**
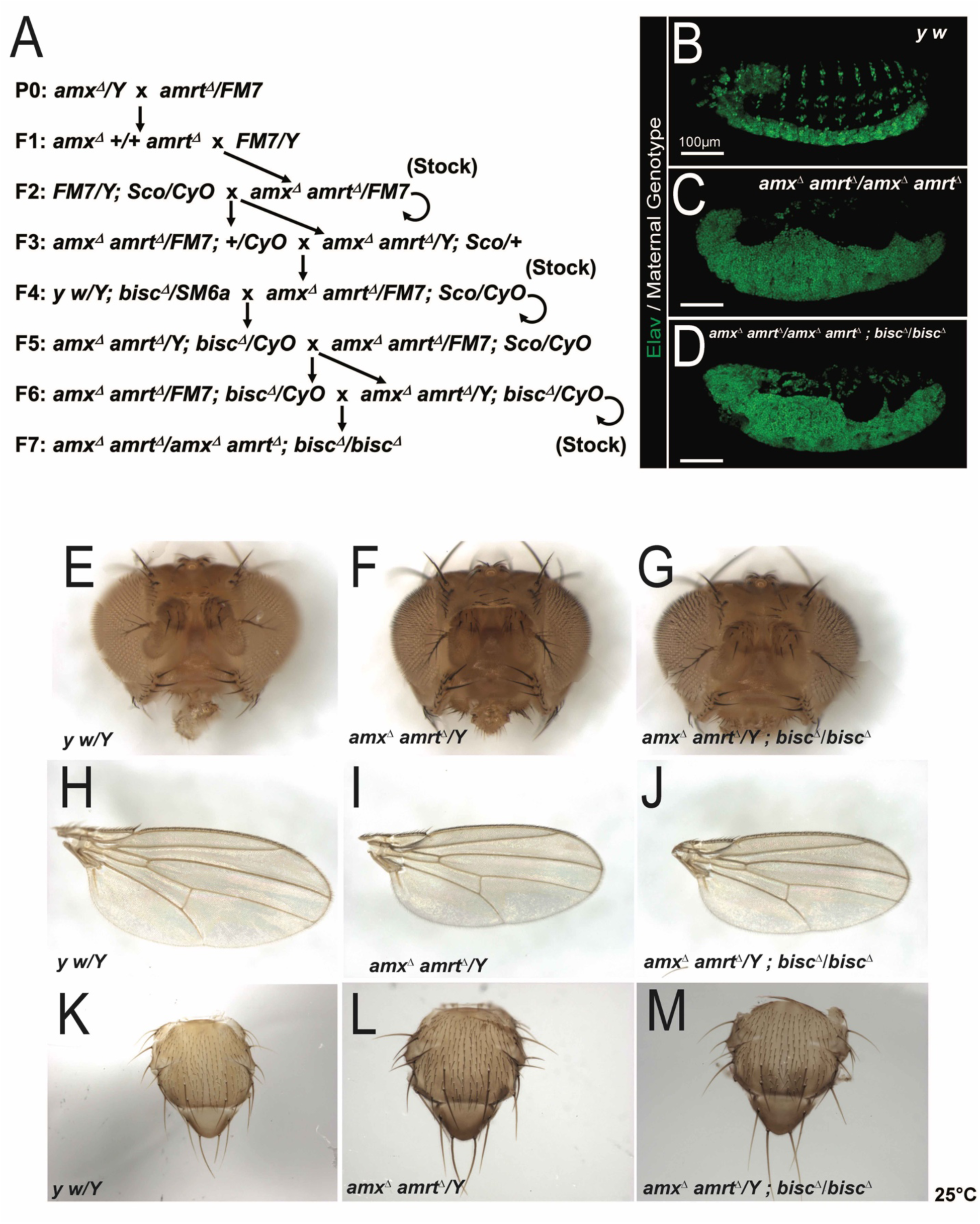
Triple null mutant for all three *TM2D* fly genes is phenotypically similar to single null mutants. (A) Crossing scheme used to generate *TM2D* triple null mutant flies. (B) A normal embryonic nervous system highlighted by neuronal nuclei marker Elav (green). (C-D) *amx^Δ^ amrt^Δ^* double mutants (C) and *amx^Δ^ amrt*^Δ^ *bisc^Δ^* triple mutants (D) exhibit a neurogenic phenotype. (E-K) *TM2D* double and triple null mutants exhibit no overt morphological phenotypes. Head structures of mutants (F, G) appear normal compared to *y w* control (E). Mutant wings (I, J) and thorax (L, M) also appear normal compared to control (H, K).

Because *amx* and *amrt* are both located on the X-chromosome (located in cytological regions 8D2 and 12C4, respectively, which are 19cM away), we recombined the two null alleles by following the dominant wing color and body color markers knocked into each locus **(Figure 3A)**. The *amx*, *amrt* double null flies (*amx^Δ^ amrt^Δ^)* in a *yellow* mutant background exhibit wild-type color wings (*y^wing2+^* marker of *amx^Δ^*) and bodies (*y^body+^* marker of *amrt^Δ^*). Absence of both *amx* and *amrt* transcripts in this line was verified by RT-PCR **(Supplemental Figure 2**). These double null adult flies also appeared morphologically normal similar to the single null animals (**Figure 3F, I, L**), suggesting that these two genes do not play redundant roles during the development of imaginal tissues. Consistent with single null animals, the *amx, amrt* double null females were fully sterile, exhibiting a maternal-effect neurogenic phenotype **(Figure 3C)**.

We next combined the *bisc^Δ^* allele on the second chromosome with the *amx^Δ^ amrt^Δ^* mutant X-chromosome to generate a triple null mutant line (*amx^Δ^ amrt^Δ^; bisc^Δ^*) (**Figure 3A**). This genetic manipulation still did not produce any adult animals with obvious Notch signaling related external morphological defects (**Figure 3G, J, M**), suggesting that these three genes also do not play redundant roles in these contexts. Consistent with the single and double null lines, the triple null mutant females are completely sterile and their progeny show maternal-effect neurogenic defects (**Figure 3D**). We did not observe any differences in the severity of the neurogenic phenotype in the embryos from the single gene knockouts and the triple knockout mothers (**Figure 1D, Figure 2C, 2G, Figure 3D**), likely because the neurogenic defect is already strong in single gene knockout animals (**Figures 1, 2**), precluding any additive or synergistic effects.

### A truncated form of Amx acts as a potent inhibitor of Notch signaling

Knock-out experiments revealed that while each of the three *TM2D* genes are individually maternally necessary for proper Notch signaling during embryogenesis, they appear to be dispensable for other developmental contexts that depend on Notch. To further study the function of TM2D proteins *in vivo*, we tested whether overexpression of Amx is sufficient to modulate Notch signaling (**Figure 4**). In addition to generating a transgene that allows the expression of a full-length Amx protein tagged with an N’-3xHA tag (placed immediately after the predicted signal sequence, 3xHA::Amx^FL^) (**Figure 4A**) using the GAL4/UAS binary expression system (Brand and Perrimon, 1993), we generated a version of this construct that lacks the majority of the non-conserved N’-terminal extracellular domain (3xHA::Amx^ΔECD^) (**Figure 4D**) to specifically test the function of the highly conserved TM2 domain.

**Figure 4.**
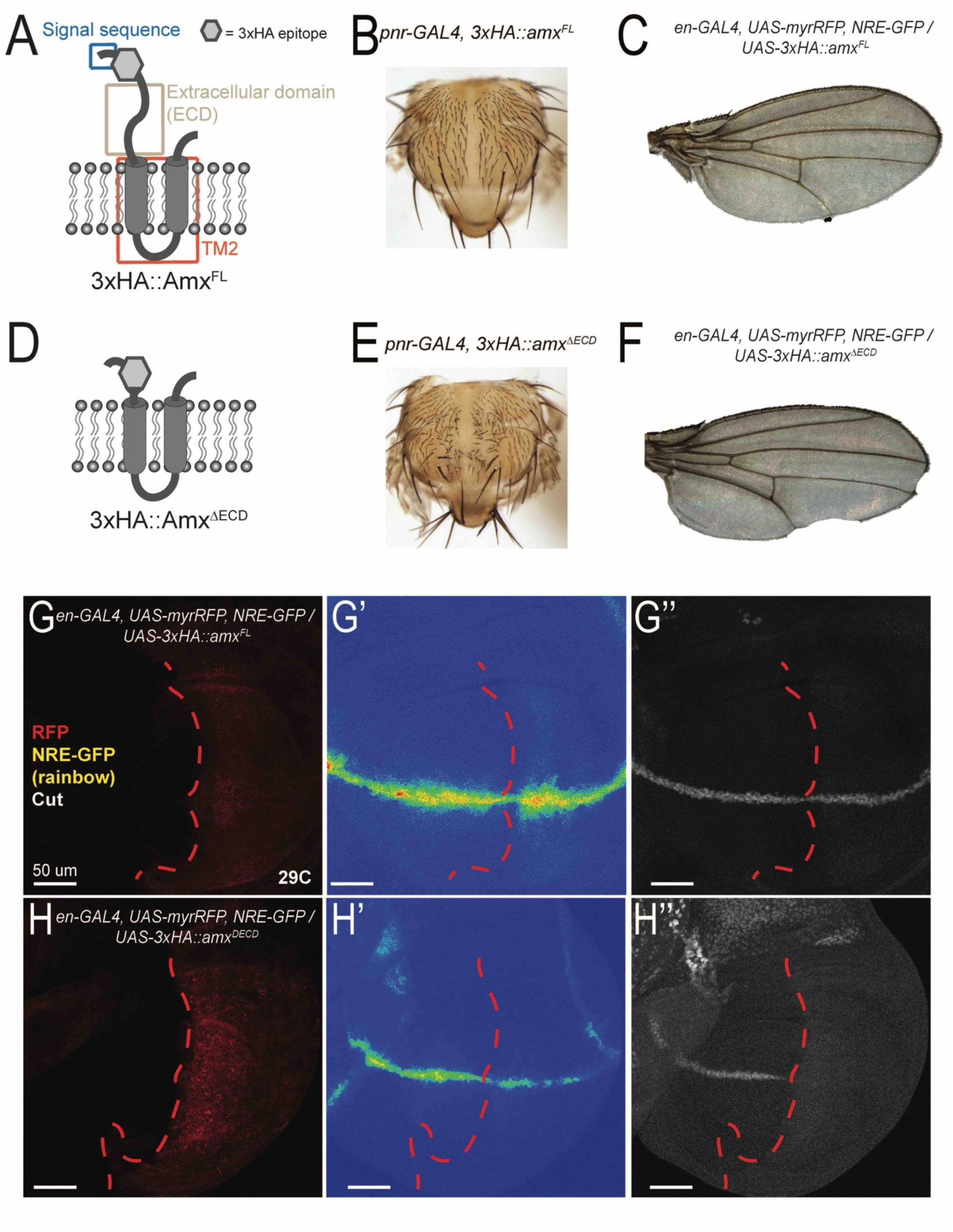
Amx that only possesses the highly conserved TM2 domain is a potent inhibitor of Notch signaling. (A, D) Schematic of proteins generated from *UAS-3xHA::amx^FL^* and *UAS-3xHA::amx^ΔECD^* transgenes. A 3xHA epitope (grey hexagon) was inserted after a predicted signal sequence (SS, blue box). The majority of the extracellular domain (ECD, light brown box) was removed to generate 3xHA::amx^ΔECD^, consisting mostly of the TM2 domain (red box) tagged with the N-terminal 3xHA epitope. (B) Overexpression of 3xHA::Amx^FL^ with *pannier(pnr)*-GAL4 has no effect on notum morphology. (C) Expression of 3xHA::Amx^FL^ in the posterior wing using *engrailed* (*en*)-GAL4 has no effect of the morphology of wings of adults raised at 29⁰C. (E) *pnr-GAL4* driven overexpression of truncated Amx causes an increase in the number of micro- and macrochaete, indicative of loss of Notch mediated lateral inhibition. (F) *en-GAL4* driven overexpression of 3xHA::Amx^ΔECD^ causes notching of the posterior wing margin. (G-H) Immunostaining of wing imaginal discs expressing full-length or truncated 3xHA::Amx. (G) 3xHA::Amx^FL^ expression in the posterior imaginal wing disc using *en-GAL4* has no effect on NRE (Notch response element)-GFP expression, a synthetic *in vivo* Notch signaling reporter (G’, rainbow) and on Cut (G’’, white) expression, a downstream target of Notch activation in this context. The domain expressing GAL4 is marked by RFP (red). (H) Expression of 3xHA::Amx^ΔECD^ decreases NRE-GFP (rainbow) expression (H’) and reduces Cut expression (H’’).

When we expressed these two transgenes in the developing dorsal thorax using *pannier-GAL4* (*pnr-GAL4*), we observed that Amx^FL^ did not cause any defects whereas Amx^ΔECD^ caused an increase in the number of mechanosensory bristles (**Figure 4B, E**), suggesting an effect on Notch mediated lateral inhibition. We then expressed the two proteins in the developing posterior compartment within wing imaginal discs using *engrailed-GAL4* (*en-GAL4*). We observed notching of the posterior wing margin when Amx^ΔECD^ was expressed (**Figure 4F**), while no such defect was seen upon expression of Amx^FL^ (**Figure 4C**). To determine whether the wing notching caused by Amx^ΔECD^ overexpression was indeed due to loss of Notch signaling, we visualized Notch activation using NRE-GFP (Notch Responsive Element-Green Fluorescent Protein), a synthetic *in vivo* Notch signaling reporter (Housden et al., 2012), as well as immunostaining of Cut, encoded by an endogenous downstream target gene of Notch activation in this context (Micchelli et al., 1997). While overexpression of Amx^FL^ did not affect NRE-GFP and Cut expression (**Figure 4G**), expression of Amx^ΔECD^ caused a reduction in both NRE-GFP and Cut expression within the *en-GAL4* expression domain (**Figure 4H**). In summary, overexpression of full-length Amx did not affect developmental events related to Notch signaling in the thorax and wing, while overexpression of a truncated form that only carries the conserved TM2 domain and its short C’-tail inhibited Notch signaling in several developmental contexts.

### Truncated Amx inhibits Notch signaling at the γ-secretase mediated receptor cleavage step

Amx has been proposed to function at the γ-secretase cleavage step of Notch activation based on a genetic epistasis experiment (Michellod and Randsholt, 2008). Notch signaling activation is initiated by the binding of the Notch receptor to its ligands (Delta or Serrate in *Drosophila*) (Hori et al., 2013). This induces a conformational change of Notch to reveal a cleavage site that is recognized by ADAM10 (encoded by *kuzbanian* in *Drosophila*) (Duojia and Rubin, 1997). Notch receptor that has undergone ADAM10 cleavage (S2 cleavage) is referred to as N^EXT^ (Notch extracellular truncation) and becomes a substrate for γ-secretase (Mumm et al., 2000). N^EXT^ that is cleaved by γ-secretase (S3 cleavage) releases its intracellular domain (N^ICD^), which then translocates to the nucleus and regulates transcription of downstream target genes (De Strooper et al., 1999). To determine how *amx* regulates Notch signaling, Michellod and Randsholt attempted to suppress the embryonic neurogenic phenotype of embryos produced from *amx^1^* mutant mothers by zygotically overexpressing different forms of Notch using a heat-shock promoter (Michellod and Randsholt, 2008). While N^ICD^ was able to weakly suppress the neurogenic defect, N^EXT^ was not able to do so, suggesting that Amx somehow modulates the function of the γ-secretase complex. However, because the phenotypic suppression observed by N^ICD^ in this study was very mild and since the authors used Notch transgenes that were inserted into different regions of the genome (thus *N^ICD^* and *N^EXT^* may be expressed at different levels and cannot be directly compared), additional data is required to fully support this conclusion.

To determine how Amx^ΔECD^ inhibits Notch signaling when ectopically overexpressed, we performed similar epistasis experiments but with improved genetic tools (**Figure 5**). First, we generated several UAS constructs expressing different forms of Notch and inserted them into the identical genomic location on the 2^nd^ chromosome using site specific ϕC31-mediated transgenesis to avoid positional effects (Bischof et al., 2012; Venken et al., 2006) (**Figure 5A**). In addition to transgenes that allow expression of N^ICD^, N^EXT^ and full-length Notch (N^FL^), we also generated a ligand-independent form of *UAS-Notch* that still depends on ADAM10 and γ-secretase by deleting several epidermal growth factor-like repeats (EGF) of the extracellular domain that contains the ligand binding domain and Lin-12/Notch Repeats (LNR) within the negative regulatory region (N^ΔEGF1-18.LNR^) (Lieber et al., 2002). When we overexpressed N^ICD^, N^EXT^ or N^ΔEGF1-18.LNR^ in the developing wing pouch using *nubbin-GAL4* (*nub-GAL4*), we observed increased Cut expression throughout the wing pouch, indicating ectopic Notch activation (**Figure 5B-D**). Over-expression of N^FL^ only showed a mild increase in Cut expression in limited regions of the wing pouch (**Supplemental Figure 3B**), likely due to its ligand-dependence. These observations are consistent with previous reports using UAS-Notch transgenic lines generated using random (P-element mediated) transgenesis technology (Doherty et al., 1996).

**Figure 5.**
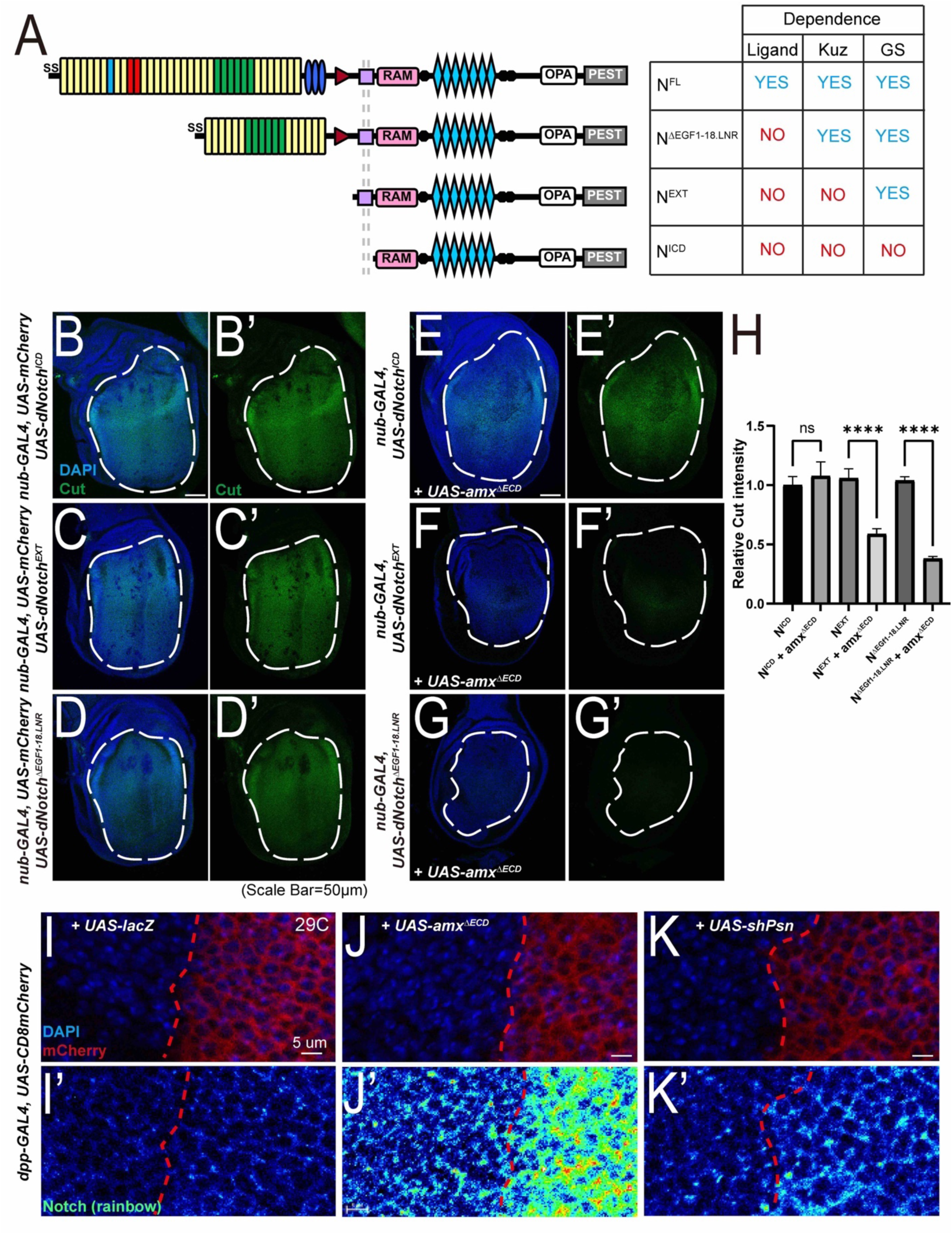
Genetic epistasis experiments place truncated Amx at the γ-secretase cleavage step of Notch activation. (A) Schematics and characteristics of Notch proteins made from each *UAS-Notch* transgenes that were generated for this study. Full-length Notch (N^FL^) requires ligand binding and processing by Kuzbanian (Kuz) and γ-secretase (GS) for activation. Notch with EGF repeats and LNR domains removed (N^ΔEGF1-18.LNR^) is not dependent on ligands but are dependent on both Kuz and GS for activation. Notch with an extracellular truncation (N^EXT^) is dependent only on GS for activation. The Notch intracellular domain (N^ICD^) is constitutively active. (B-D) Expression of Notch constructs leads to increase in Cut (green) expression, quantified in (H). (E) Co-overexpression of 3xHA::Amx^ΔECD^ has no effect of N^ICD^ mediated increases in Cut expression (E’, H). (F-G’) 3xHA::Amx^ΔECD^ expression suppresses the effects of N^ΔEGF1- 18.LNR^ and N^EXT^ on Cut expression (F’, G’, H). t-test. *= p<0.05. ****= p<0.0001. Error bars show SEM. Scale bar = 50 µm. (I-K) Overexpression of 3xHA::Amx^ΔECD^ causes an increase of Notch protein levels (J’) compared to overexpression of a neutral protein, LacZ (β-galactosidase) (I’). Knockdown of *psn* mediated by shRNA also results in mild increase Notch levels (K’), mimicking the effect of 3xHA::Amx^ΔECD^.

Next, we recombined *nub-GAL4* and *UAS-3xHA::amx^ΔECD^* onto the same chromosome. The resultant genetic recombinants constitutively overexpresses Amx^ΔECD^ in the wing pouch, with heterozygous adults exhibiting wing notching and homozygous animals completely lacking wings in adulthood (**Supplemental Figure 4A-B**). We then crossed these *nub-GAL4, UAS-3xHA::amx^ΔECD^* flies to different *UAS-Notch* lines to determine if Amx^ΔECD^ can modulate the ectopic Cut expression phenotype caused by Notch overexpression. We found that co-overexpression of Amx^ΔECD^ significantly suppresses the induction in Cut expression caused by N^ΔEGF1-18.LNR^ or N^EXT^ (**Figure 5E-F**), but had no effect on the same phenotype caused by N^ICD^ (**Figure 5G**). This indicates that *amx^ΔECD^* genetically acts at the γ-secretase-mediated cleavage step of Notch activation, consistent with previous epistasis experiments using *amx^1^* (Michellod and Randsholt, 2008).

To further understand how Amx^ΔECD^ inhibits Notch signaling upon overexpression, we assessed the distribution of the Notch receptor through immunostaining using a monoclonal antibody that recognizes both cleaved and uncleaved forms of Notch (Fehon et al., 1990). Upon overexpression with *decapentaplegic-GAL4* (*dpp-GAL4*), which is expressed in a limited domain within the wing pouch, we observed that Amx^ΔECD^ causes wing notching (**Supplemental Figure 4D**) that is accompanied by a dramatic upregulation of Notch receptor (**Figure 5J-K**). To test whether a similar phenotype is seen upon loss of γ-secretase function, we knocked-down *Psn (Presenilin*, which is orthologous to human *PSEN1* and *PSEN2)* in the wing disc and assessed its effect on Notch protein levels. When we performed RNAi against *Psn* using a *UAS-RNAi* line that had been validated in a previous study (Kang et al., 2017) with *dpp-GAL4*, we found that knock-down of *Psn* shows an accumulation of Notch similar to Amx^ΔECD^ over-expression but to a lesser extent (**Figure 5L**). To also test whether this Notch accumulation phenotype occurs when Notch cleavage is altered by another mechanism, we tested whether loss of ADAM10 also causes this defect. When we generated mutant clones of a null allele of *kuz* (*kuz^e29-4^*) (Rooke et al., 1996) using the MARCM (Mosaic analysis with a repressible cell marker) system (Lee and Luo, 1999), we did not observe any alterations in Notch expression (**Supplemental Figure 5**), indicating that the increase in Notch levels seen upon over-expression of Amx^ΔECD^ and *Psn* knockdown is rather specific.

In summary, while over-expression of the full-length Amx protein did not cause any obvious defects, we serendipitously found that a truncated form of this protein that only contains the region that is highly conserved among all TM2D proteins can act as a potent inhibitor of Notch signaling when overexpressed. Epistasis experiments show that Amx^ΔECD^ acts at the S3 cleavage step of Notch activation, suggesting that it likely regulates γ-secretase. This is further supported by our findings that overexpression of Amx^ΔECD^ and knock-down of *Psn*, but not loss of *kuz*, both lead to accumulation of Notch.

### *amx* null mutants have shortened lifespan

AD is an adult-onset age-dependent disease that worsens over time. While *TM2D3^p.P155L^* has been associated with LOAD (Jakobsdottir et al., 2016) and *TM2D3^p.P69L^* has been recently reported in a proband with EOAD or frontotemporal dementia (Cochran et al., 2019), there is no functional data that directly links this gene to an age-dependent neurological phenotype in any species. To determine if loss of *TM2D3* causes an age-dependent phenotype, we first assessed the lifespan of *amx^Δ^* mutant animals. We compared the longevity of *amx^Δ^* hemizygous male flies with flies that also carry a genomic rescue construct (*amx^Δ^ + amx*) to minimize the effect of genetic backgroud (Jakobsdottir et al., 2016). We selected males for our analysis because *amx* loss does not affect male fertility, allowing us to ignore any changes in lifespan that may be caused by alterations in fecundity (Flatt, 2011). In contrast to the rescued control animals that exhibit a median lifespan of 51 days, the median lifespan of *amx* null flies is significantly shorter at 27 days (**Figure 6A**, p=1.0×10^-11^). We next tested whether human *TM2D3* can substitute for the loss of *amx* in this context by introducing the humanized genomic rescue construct (Jakobsdottir et al., 2016) into the *amx* null mutant background (*amx^Δ^ + hTM2D3*). These humanized *TM2D3* animals had a median lifespan of 33 days (**Figure 6A**). These data indicate that loss of *amx* causes reduction in lifespan, and human *TM2D3* can weakly but significantly (p=9.5×10^-10^) suppress this phenotype.

**Figure 6.**
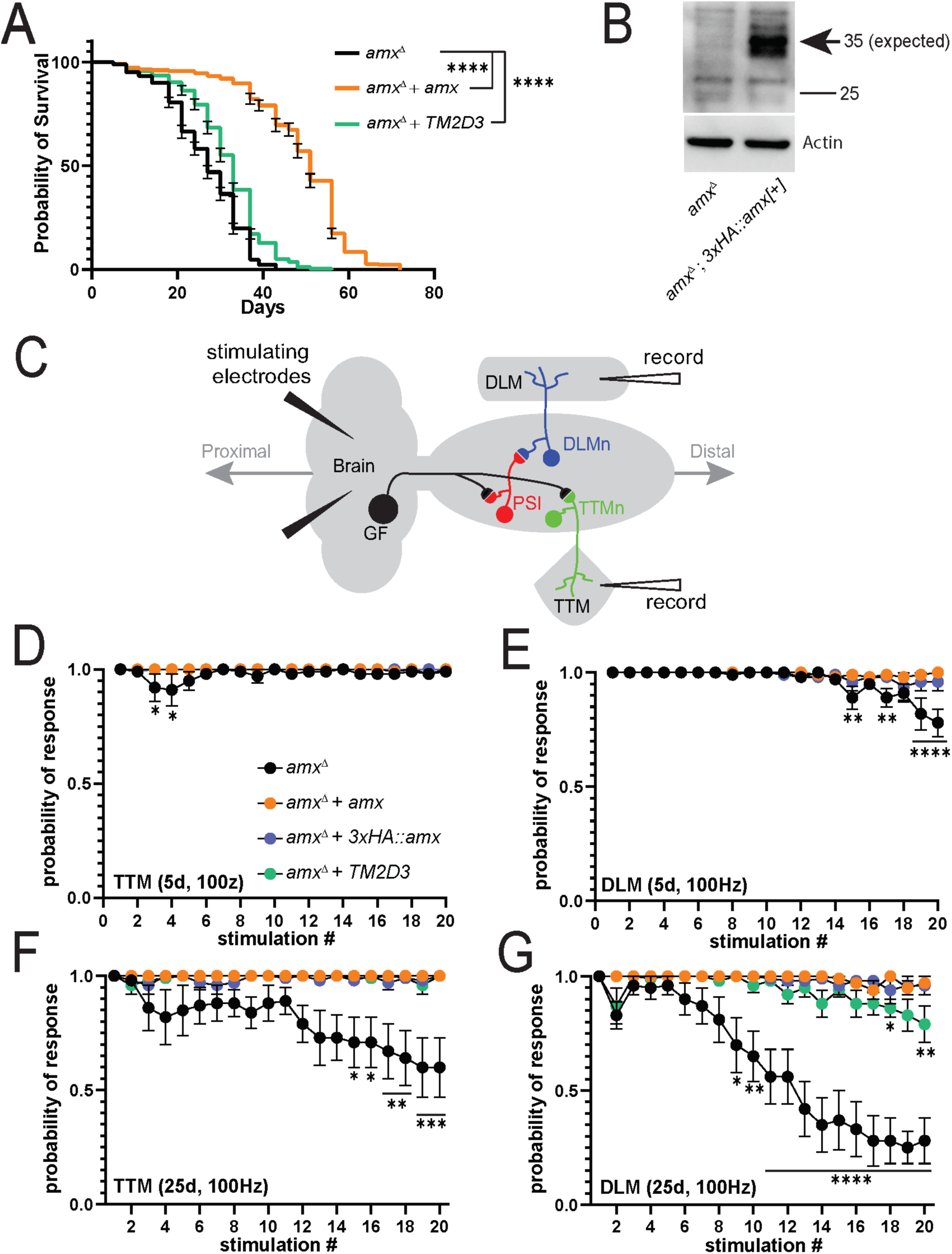
Loss of *amx* causes shortening of lifespan and age-dependent neurophysiological defects. (A) Lifespan assay shows *amx^Δ^* animals (black, n=247) have reduced lifespan compared to *amx^Δ^ + amx* controls (orange, n=224). *amx^Δ^ +* human *TM2D3* flies (green, n=234) have significantly longer lifespan than *amx^Δ^* animals, but shorter than control. Animals were reared at 25⁰C; Log-rank test (Mantel-Cox), ****= p<0.0001. (B) Western blot of *amx^Δ^; 3xHA::amx* brains shows that 3xHA::Amx (predicted 35 kDa size) is expressed in the adult nervous system (arrow). Protein isolate from five brains was loaded per lane and Actin was probed as a loading control. (C) Schematization of the giant fiber electrophysiological recordings. Stimulating electrodes are inserted into the brain and recording electrodes record responses from the TTM and DLM muscles. (D-E) TTM muscles of 5d old *amx^Δ^* mutants (black) have a response similar to *amx^Δ^ + amx* controls (orange) while DLM muscles have small but significant decrease in response probability. *3xHA::amx* (blue) flies also perform as well as controls. (F-G) TTM and DLM response in 25d old *amx^Δ^* mutants is significantly reduced. DLM response of 25d old *amx^Δ^ +* human *TM2D3* (green) flies is reduced compared to controls (I). Multiple unpaired t-tests with Holm-Šídák correction for multiple comparisons. *= p<0.05. ** = p≤0.01. ***= p<0.001, ****= p<0.0001. Error bars show SEM. Additional data can be found in **Supplemental Figures 8-10**.

### Amx is expressed in the adult brain

Shortened lifespan seen in *amx^Δ^* animals could be due to a number of reasons including defects in the nervous system or other organs functionally affected by the loss of *amx*. Based on publicly available microarray (Chintapalli et al., 2007), RNA sequencing (RNA-seq) (Brown et al., 2014) and single cell RNA-sequencing (scRNA-seq) (Davie et al., 2018) datasets, *amx* mRNA has been detected in the adult nervous system at low levels in a subset of neurons and glia cells, though it’s expression level in the nervous system is higher than most other tissues examined. (**Supplemental Figure 6**). To determine whether Amx protein can be detected in the adult nervous system, we generated an N’-tagged *amx* genomic rescue construct in which a 3xHA epitope is inserted immediately after the signal sequence of the Amx protein (**Figure 1A-B**) similar to the *UAS-3xHA::amx* transgene (**Figure 4A**). This tagged genomic rescue construct is able to rescue the female fertility and the maternal-effect neurogenic phenotype of *amx^Δ^* (**Figure 1C, 1F**), indicating the epitope tag does not have a major effect on Amx function. Next, we verified expression of 3xHA::Amx in the female ovary through immunofluorescent staining and western blot (**Supplemental Figure 7**). Based on immunostaining, we detected 3xHA::Amx in nurse cells of the ovary and observed that it localizes to the cell membrane as well as in intracellular puncta (**Supplemental Figure 7A-B**). Based on western blot, we identified 3xHA::Amx at the predicted molecular weight [34.82 kDa (31.35 kDa for Amx, 3.47 kDa for the 3xHA tag)] (**Supplemental Figure 7C**). Finally, we assessed the expression of 3xHA::Amx in the adult brain via immunostaining and western blot. While we did not observe a strong signal beyond background fluorescence based on immunofluorescence staining using an anti-HA antibody (not shown), we detected 3xHA::Amx via western blot using brain extracts at the expected molecular weight as we observed in the ovary extracts (**Figure 6C**). In conclusion, Amx is expressed in the adult nervous system at relatively low levels.

### *amx* null mutants show progressive electrophysiological defects

Finally, to directly assess whether *amx* is required for maintenance of neural function, we performed electrophysiology on *amx^Δ^* animals and controls. The giant fiber system is a neuronal circuitry that is required for rapid escape responses in insects and has been used as a model circuitry to assess neuronal function in a quantitative manner in adult flies (Allen and Godenschwege, 2010). This pathway can be activated through direct stimulation of the brain and the outputs of the circuit can monitored by recording the responses from the tergotrochanteral (TTM) and dorsal longitudinal (DLM) muscles (**Figure 6C**). For a given time point, we applied multiple stimulations at different frequencies (20, 50, and 100 Hz; **Figure 6D-G**, **Supplemental Figures 8-10**), and measured how well these muscles respond to each stimulation. Healthy neurons can follow the stimulations showing a ‘probability of response’ that is close to 1.0, but neurons that are unhealthy show decreased ‘probability of response’, which indicates failure of the signal to travel from the brain to the muscles (Martinez et al., 2007; Oyston et al., 2018).

At 5 days post-eclosion, *amx^Δ^* flies have a minor but significant failure rate at later stimulations at 100 Hz in the DLM muscle compared to control animals (rescued with a wild-type *amx* [*amx^Δ^ + amx*] or N’-3xHA tagged *amx* [*amx^Δ^ + 3xHA::amx*]; **Figure 6E**). Additionally, *amx^Δ^* animals show only slight, early stimulation failure rates in the TTM at 50 and 100 Hz at 5 days (**Figure 6D, Supplemental Figure 8D**). At 15 days post-eclosion, *amx^Δ^* animals still only show slight failure rate in the TTM at 100 Hz (**Supplemental Figure 9E**) but begin to show a significant increase in failure rate compared to control animal recordings from the DLM at 50 and 100 Hz (**Supplemental Figure 9D, F**). At 25 days post-eclosion, the defects in the DLM increase in severity for 50 and 100 Hz compared to rescued controls (**Figure 6G, Supplemental Figure 10D**). At this time point, the TTM also begins to show a significant failure rate in *amx^Δ^* animals compared to rescued animals (**Figure 6F**). Interestingly, unlike in the lifespan assay in which the human TM2D3 did not dramatically suppress the *amx^Δ^* mutant phenotype and in the fertility/neurogenesis assay in which the human TM2D3 showed ∼50% activity of the fly Amx protein, we found that human TM2D3 is able to significantly suppress the electrophysiological defects close to the level of fly Amx (**Figure 6F-G**). In summary, loss of *amx* causes an age-dependent decline in neuronal function, and this defect can be fully rescued by both fly and human *TM2D3*, indicating an evolutionarily conserved role of this gene in healthy aging.

## Discussion

In this study, we functionally characterized *TM2D* genes through gene knockout and over-expression strategies in *Drosophila melanogaster* to gain biological knowledge on this understudied but evolutionarily conserved gene family that has been implicated in AD. We first showed that the knockout allele of *amx* (*Drosophila* homolog of *TM2D3*) generated by CRISPR is phenotypically indistingusiable from the classic *amx^1^* allele and displays female sterility and a maternal-effect neurogenic defect. Recently, we reported that this allele also shows a maternal-effect inductive signaling defect to specify the mesoectoderm during embryogenesis, which is another Notch-dependent event (Das et al., 2020), demonstrating that *amx* is maternally required for multiple Notch signaling dependent processes during embryogenesis. In addition, we generated the first knockout alleles of *amrt* (*Drosophila* ortholog of *TM2D2*) and *bisc* (*Drosophila* ortholog of *TM2D1*) and documented that each null allele phenotypically mimics the loss of *amx*. Furthermore, we revealed that the triple knockout of all three *TM2D* genes in *Drosophila* show identical maternal-effect neurogenic phenotypes without exhibiting other obvious Notch signaling-related developmental defects. Moreover, although the over-expression of the full-length Amx did not cause any scorable defects, we serendipitously found that expression of a truncated form of Amx that lacks the majority of the extracellular domain (Amx^ΔECD^) can strongly inhibit Notch signaling in the developing wing imaginal disc. Through genetic epistatic experiments using newly generated *UAS-Notch* transgenic lines, we mapped this inhibitory effect to the γ-secretase cleavage step of Notch activation. Subsequently, we showed that Amx is expressed in the adult nervous system and that *amx* null animals have a shortened lifespan. Finally, through electrophysiological recordings of the giant fiber system, we showed that *amx* null flies show age-dependent decline in neuronal function. In summary, we demonstrate that all three *TM2D* genes play critical roles in embryonic Notch signaling to inhibit the epithelial-to-neuron cell fate transformation as maternal-effect genes, and that *amx* is required for neuronal maintenance in the adult nervous system, a function that may be related to the role of human *TM2D3* in AD.

*TM2D* genes are found in multicellular animals but are absent in yeasts (e.g. *Saccharomyces cerevisiae*, *Schizosaccharomyces pombe*) and plants (e.g. *Arabidopsis thaliana*), suggesting that this family of genes arose early in the metazoan lineage. In humans and flies, there are three *TM2 domain-containing* genes (*TM2D1*, *TM2D2*, *TM2D3* in *Homo sapiens*; *bisc*, *amrt*, *amx* in *Drosophila melanogaster*, respectively), each corresponding to a single gene in the other species. Interestingly, this 1:1 ortholog relationship is also seen in mouse (*Tm2d1*, *Tm2d2*, *Tm2d3*), frog (*Xenopus tropicalis*: *tm2d1*, *tmd2d*, *tm2d3*) zebrafish (*Danio rerio*: *tm2d1*, *tm2d2*, *tm2d3*) and worm (*Caenorhabditis elegans*: *Y66D12A.21*, *C02F5.13*, *C41D11.9*) (**Supplemental Figure 1**). In general, most genes have more paralogous genes in humans compared to flies (for example, one *Drosophila Notch* gene corresponding to four *NOTCH* genes in human) as vertebrates underwent two rounds of whole-genome duplication (WGD) events during evolution (Kasahara, 2007). Furthermore, teleosts including zebrafish underwent an extra round of WGD (Glasauer and Neuhauss, 2014), leading to formation of extra duplicates in 25% of all genes (e.g. one *NOTCH1* gene in human corresponds to *notch1a* and *notch1b* in zebrafish). Hence it is interesting that each of the three *TM2D* genes remained as single copy genes in various species despite whole genome level evolutional changes, suggesting that there may have been some selective pressure to keep the dosage of these genes consistent and balanced during evolution.

Although the *in vivo* functions of *TM2D1* and *TM2D2* have not been studied in any organism, several lines of studies performed in cultured cells suggest that these genes may also play a role in AD pathogenesis. Through a yeast-two hybrid screen to identify proteins that bind to Aβ42, Kajkowski et al. identified TM2D1 and referred to this protein as BBP (beta-amyloid binding protein) in their study (Kajkowski et al., 2001). They further showed that TM2D1 can also interact with Aβ40, a non-amyloidogenic form of Aβ, and mentioned that they have preliminary data that it also binds to APP (published and unpublished data in Kajkowski et al., 2001). The interaction between Aβ peptides and TM2D1 was shown to require the extracellular domain as well as a portion of the first transmembrane domain of TM2D1. Because overexpression of TM2D1 in a human neuroblastoma cell line (SH-SY5Y) increased the sensitivity of these cells to cell death caused by incubation with aggregated Aβ and since the DRF motif was found to be required for this activity, the authors of this original study proposed that TM2D1 may function as a transmembrane receptor that mediates Aβ-toxicity (Kajkowski et al., 2001). However, a follow-up study from another group refuted this hypothesis by providing data that TM2D1 is not coupled to G proteins using a heterologous expression system in *Xenopus* oocytes (Lee et al., 2003).

To our surprise, loss of *bisc/TM2D1* and *amrt/TM2D2* were phenotypically indistinguishable from the loss of *amx/TM2D3*. The zygotic loss of each gene did not exhibit any strong developmental defects into adulthood, despite their relatively ubiquitous expression pattern according to large transcriptome datasets (Brown et al., 2014; Chintapalli et al., 2007). Similarly, the triple null mutants did not exhibit any morphological defect, suggesting that these genes are not required zygotically during development. In contrast, maternal loss of any single *TM2D* gene causes a strong neurogenic defect, which is also seen in embryos laid by triple knockout animals. The neurogenic defect is a classical phenotype in *Drosophila* that was originally reported in the mid-1930s (Artavanis-Tsakonas and Muskavitch, 2010; Yamamoto et al., 2014), and the study of mutants that show this phenotype led to the establishment of the core Notch signaling pathway in the late 1980’s and early 1990’s (Artavanis-Tsakonas et al., 1995; Lehmann et al., 1981). Although the study of neurogenic phenotypes and genes has a long history, this phenotype is a very rare defect that has so far been associated with only 18 genes according to FlyBase (Larkin et al., 2021), prior to this work. Seven genes show this defect as zygotic mutants [*aqz*, *bib*, *Dl*, *E(spl)m8-HLH*, *mam*, *N* and *neur*], seven genes are zygotically-required essential genes with large maternal contributions (hence the need to generate maternal-zygotic mutants by generating germline clones to reveal the embryonic neurogenic defect) [*Gmd*, *Gmer*, *gro*, *Nct*, *O-fut1*, *Psn* and *Su(H)*], one gene has only been investigated by RNAi (*Par-1*) and four genes including *amx* are non-essential genes and show maternal-effect neurogenic defects (*amx*, *brn*, *egh*, *pcx*). Hence, our study has revealed two new genes that are evolutionarily closely linked to *amx* in this Notch signaling related process.

The similarity of the phenotypes and sequences of *amx*, *amrt* and *bisc* suggests that these proteins may function together in the context of embryonic neurogenesis. Interestingly, high-throughput proteomics data based on co-immunoprecipitation mass spectrometry (co-IP/MS) from human cells has detected physical interactions between TM2D1-TM2D3 (Oughtred et al., 2021) and TM2D2-TM2D3 (Huttlin et al., 2017), suggesting these proteins may form a protein complex. Further biochemical studies will be required to clarify the functional relationship between the three TM2D proteins. Two additional mammalian datasets further support our hypothesis that these three proteins functions together. First, all three *TM2D* genes were identified through a large scale cell-based CRISPR-based screen to identify novel regulators of phagocytosis (Haney et al., 2018). Singular knock-out of *TM2D* genes in a myeloid cell line was sufficient to cause a similar phagocytic defect among the mutant cell lines based on the parameters the authors screened for (e.g. substrate size, materials to be engulfed). Although the authors of this study did not generate double or triple knockout cell lines to determine whether there were additive or synergistic effects when multiple *TM2D* genes were knocked out, this suggests that the three genes may function together in phagocytosis. The authors further note these genes are broadly expressed in diverse cell types beyond phagocytic cells in the nervous system and thus may play other roles in disease progression (Zhang et al., 2014). Second, preliminary phenotypic data from the International Mouse Phenotyping Consortium (Dickinson et al., 2016) indicates that single knockout of mice of *Tm2d1* (https://www.mousephenotype.org/data/genes/MGI:2137022), *Tm2d2* (https://www.mousephenotype.org/data/genes/MGI:1916992) and *Tm2d3* (https://www.mousephenotype.org/data/genes/MGI:1915884) are all recessive embryonic lethal prior to E18.5. Although detailed characterization of these mice will be required and further generation of a triple knockout line is desired, the shared embryonic lethality may indicate that these three genes potentially function together in an essential developmental paradigm during embryogenesis in mice.

Our attempts to unravel the function of Amx through overexpression of the full-length protein was uninformative since this manipulation did not cause any scorable phenotype. However, we found that expression of the most conserved region of Amx that lacks the majority of the N’-extracellular domain strongly inhibited Notch signaling during wing and notum development. These results were surprising because we did not see any wing or bristle defects in the triple *TM2D* gene family knockout flies. This could be due to one of the two following possibilities: First, *TM2D* genes do indeed play regulatory roles during wing and bristle development but zygotic mutants do not show any phenotypes because there is sufficient maternal contribution (the zygotic triple knockout *amx^Δ^ amrt^Δ^; bisc^Δ^* flies are derived from *amx^Δ^ amrt^Δ^/+ +; bisc^Δ^/+* females who have one copy of each *TM2D* gene still intact). This hypothesis is supported by high-throughput transcriptomics (microarray and RNA-seq) data that *TM2D* genes are expressed in the ovary at a higher level compared to most other tissues (Brown et al., 2014; Chintapalli et al., 2007). In this case, Amx^ΔECD^ may be acting as a dominant-negative protein, sequestering the endogenous substrates of Amx (and potentially of Amrt and Bisc as well) because it likely only carries one of the critical functional domains of this protein (TM2 domain). The second possibility is that Amx^ΔECD^ acts as a neomorphic allele, inhibiting a protein that is involved in Notch signaling in which the conserved portion of Amx has the capacity of binding to. In either scenario, by performing epistasis analysis using different forms of Notch, we determined that the factor that Amx^ΔECD^ acts on is likely to be the γ-secretase complex. This data is consistent with earlier epistasis experiments performed on *amx^1^* in the context of embryonic neurogenesis (Michellod and Randsholt, 2008), further supporting the idea that *amx* has the capacity to regulate γ-secretase *in vivo*. We further determined that over-expression of Amx^ΔECD^ in the wing imaginal disc causes an accumulation of Notch protein, similar to what is seen upon knockdown of *Psn* (fly ortholog of human *PSEN1* and *PSEN2*), consistent with earlier findings showing Notch accumulation at the cell membrane in the neuroblasts of *Psn* mutants (Guo et al., 1999). In summary, we showed that ectopic over-expression of a portion of Amx that is conserved among TM2D proteins causes a strong Notch signaling defect due to a defect in γ-secretase function, suggesting that endogenous function of this protein is likely realted to γ-secretase, providing a potential molecular link to AD pathogenesis in humans.

By aging the *amx* null male flies that are visibly indistinguishable from the control flies (*amx* null flies with genomic rescue constructs), we found that loss of *amx* causes a significant decrease in lifespan. By generating a functional genomic rescue transgene in which Amx is tagged with an epitope tag, we found that this protein is expressed in the adult brain. By further performing electrophysiological recordings of the giant fiber system, which is a model circuit that is frequently used in neurological and neurodegenerative research in *Drosophila* (Allen and Godenschwege, 2010; Luan et al., 2014; Watson et al., 2008; Zhao et al., 2010), we found that there is an age-dependent decline in the integrity of this circuit. We observed that the DLM branch of the giant fiber system begins to show failures earlier than the TTM branch. The DLM is activated by giant fiber neurons that chemically synapse onto PSI (peripherally synapsing interneuron) neurons through cholinergic synapses, which in turn chemically synapse onto motor neurons (DLMn which are glutamatergic) through cholinergic connections. The TTM, in contrast, is activated by GF neurons that electrically synapse onto motor neurons (TTMn which are glutamatergic) through gap junctions, causing a more rapid response. Considering the difference in the sensitivity of the two branches, cholinergic neurons/synapses may be more sensitive to the loss of *amx*, a neuronal/synaptic subtype that is severely affected in Alzheimer’s disease in an age-dependent manner (Hampel et al., 2018).

How does *amx* maintain neuronal function in aged animals and is this molecular function related to AD? One potential molecular mechanism is through the regulation of γ-secretase in the adult brain. By knocking down subunits of the γ-secretase complex, *Psn* and *Nct (Nicastrin)*, specifically in adult neurons, Kang et al. showed that reduction of γ-secretase function decreases lifespan, which was associated with histological signs of neurodegeneration (Kang et al., 2017). The requirement of γ-secretase components in neuronal integrity has also been reported in mice (Feng et al., 2004; Saura et al., 2004; Tabuchi et al., 2009; Watanabe et al., 2014; Wines-Samuelson et al., 2010), suggesting this is an evolutionarily conserved phenomenon. Interestingly, the role of the γ-secretase complex in neuronal maintenance is unlikely to be due to defects in Notch signaling because neurodegeneration has not been observed upon conditional removal of Notch activity in post-developmental brains in flies and in mice (Salazar et al., 2020). While the precise function of γ-secretase in neuronal maintenance is still unknown, several possibilities including its role in regulating mitochondrial morphology (Wines-Samuelson et al., 2010) and calcium homeostasis (Wu et al., 2013; Zhang et al., 2009) based on studies in *C. elegans* and mice. Investigating whether Amx does indeed regulate γ-secretase in adult neurons and whether it impacts the aforementioned processes will likely facilitate our understanding on how this gene regulates neuronal health. Furthermore, considering that *TM2D3* and other *TM2D* genes have been proposed to function in phagocytic cells, and because phagocytosis process plays many roles beyond engulfment of toxic Aβ molecules in the nervous system (Fu et al., 2014), Amx may also be playing a role in engulfing unwanted materials that are harmful for the adult brain. For example, loss of the phagocytic receptor Draper in glia cells causes age-dependent neurodegeneration that is accompanied by accumulation of non-engulfed apoptotic neurons throughout the fly brain (Etchegaray et al., 2016). Interestingly, a recent study has shown that over-expression of phagocytic receptors can also promote neurodegeneration (Hakim-Mishnaevski et al., 2019), indicating the level of phagocytic activity needs to be tightly controlled *in vivo*. Further studies of *amx^Δ^* mutants (as well as *amx^Δ^ amrt^Δ^; bisc^Δ^* triple mutants) in the context of phagocytosis will likely reveal the precise molecular function of Amx and other TM2D proteins in this process.

Finally, could there be any molecular link between the role of *TM2D* genes in Notch signaling (proposed based on experiments in *Drosophila*) and phagocytosis (revealed based on mammalian cell culture based studies), or are they two independent molecular functions of the same proteins? All TM2D proteins have two transmembrane domains connected by a short intracellular loop, making them an integral membrane protein. By tagging the *amx* genomic rescue construct with a 3xHA tag that does not influence the function of Amx, we observed that 3xHA::Amx is localized to the plasma membrane as well as intracellular puncta, which likely reflects intracellular vesicles. Interestingly in embryos laid by *amx^Δ^* mutant females, we observed a mild and transient but significant alteration in Notch distribution during early embryogenesis (Das et al., 2020). Moreover, we observed a strong accumulation of Notch when we overexpressed Amx^ΔECD^ in the developing wing primordium. These data indicate that *amx* may affect protein trafficking, which in turn may impact the processing of Notch by the γ-secretase complex. Indeed, Notch signaling is highly regulated by vesicle trafficking and alterations in exocytosis, endocytosis, recycling and degradation all impact the signaling outcome (Schnute et al., 2018; Yamamoto et al., 2010). In fact, multiple studies have proposed that γ-secretase cleavage occurs most effectively in acidified endocytic vesicles (Baron, 2012; Fortini and Bilder, 2009). Hence, while *amx* may be specifically required for the proper assembly or function of the γ-secretase complex, it may alternatively be necessary to bring Notch and other substrates to the proper subcellular location for proteolytic cleavages to occur efficiently. This latter model indicates that the primary function of Amx is to regulate intracellular trafficking, which may also explain how this protein may be involved in phagocytosis. Similar to Notch signaling, phagocytosis requires coordination of many cellular trafficking events to expand the plasma membrane to form a phagophore, internalize the particle of interest to generate a phagosome, and fuse the phagosome to lysosomes to degrade its content (Melcarne et al., 2019). By studying the role of *TM2D* genes and proteins in embryonic Notch signaling, phagocytosis and age-dependent neuronal maintenance, we will likely understand the precise molecular function of this evolutionarily conserved understudied protein family, which may lead to further understanding of molecular pathogenesis of AD and other human diseases. Considering the phenotypic similarities of *amrt* and *bisc* to *amx* in *Drosophila* neurogenesis, the similarities between *TM2D1-3* in human cells in the context of phagocytosis, and the similarities of *Tm2d1-3* knockout mice in the context of embryogenesis, we propose that rare genetic variants, epigenetic regulators or proteomic changes in other *TM2D* genes may reveal novel risk factors or biomarkers in epidemiologic study of AD and other forms of dementia.

## Acknowledgements

This work was supported by the Alzheimer’s Association New Investigator Research Grant (NIRG-15-364099) and Nancy Chang, Ph.D. Award for Research Excellence to S.Y. P.C.M. is supported by the Canadian Institutes of Health Research (MFE-164712). Confocal imaging performed in this study was supported by the Eunice Kennedy Shriver Intellectual and Developmental Disabilities Research Center (IDDRC) at Baylor College of Medicine (P50-HD10355). We are grateful to Danqing Bei and Hongling Pan for *Drosophila* microinjections. We acknowledge the Knockout Mouse Phenotyping Project (KOMP2) and Internationational Mouse Phenotyping Consortium (IMPC) for the generation and phenotyping of *Tm2d1-3* knockout mouse strains. We thank Dr. Oguz Kanca for technical advice and useful suggestions. We thank Drs. Joshua Shulman, Hugo Bellen, Bart De Strooper, Katrien Horré, Motoo Kitagawa, Wataru Masuda, Kenji Matsuno and Tomoko Yamakawa for valuable discussions. We thank Shelley Gibson and J. Michael Harnish for helpful comments on the manuscript.

## Materials and Methods

### Drosophila strains and fly husbandry

*Drosophila melanogaster* stocks used in this study are listed in the **Key Resources Table**. Some strains were generated in house for this study (see below), and others were obtained from Bloomington *Drosophila* Stock Center and other sources. Flies were kept on standard media and maintained at room temperature (21-23 °C). Crosses were performed at 25 °C in an incubator unless otherwise stated.

### Genotype of Flies used in each Figure panel

The genotypes of the flies shown in each figure panel are listed in the table below:

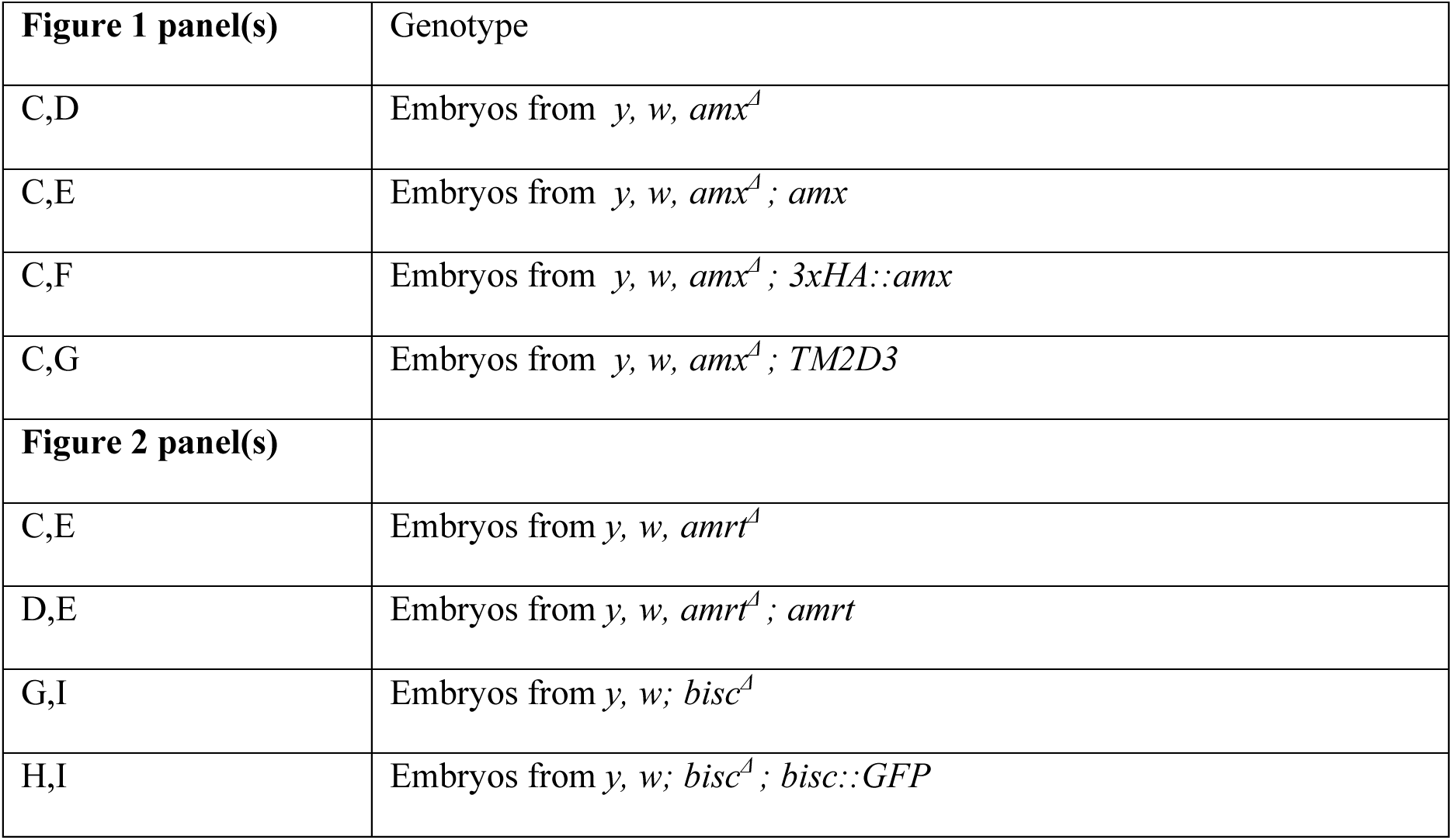
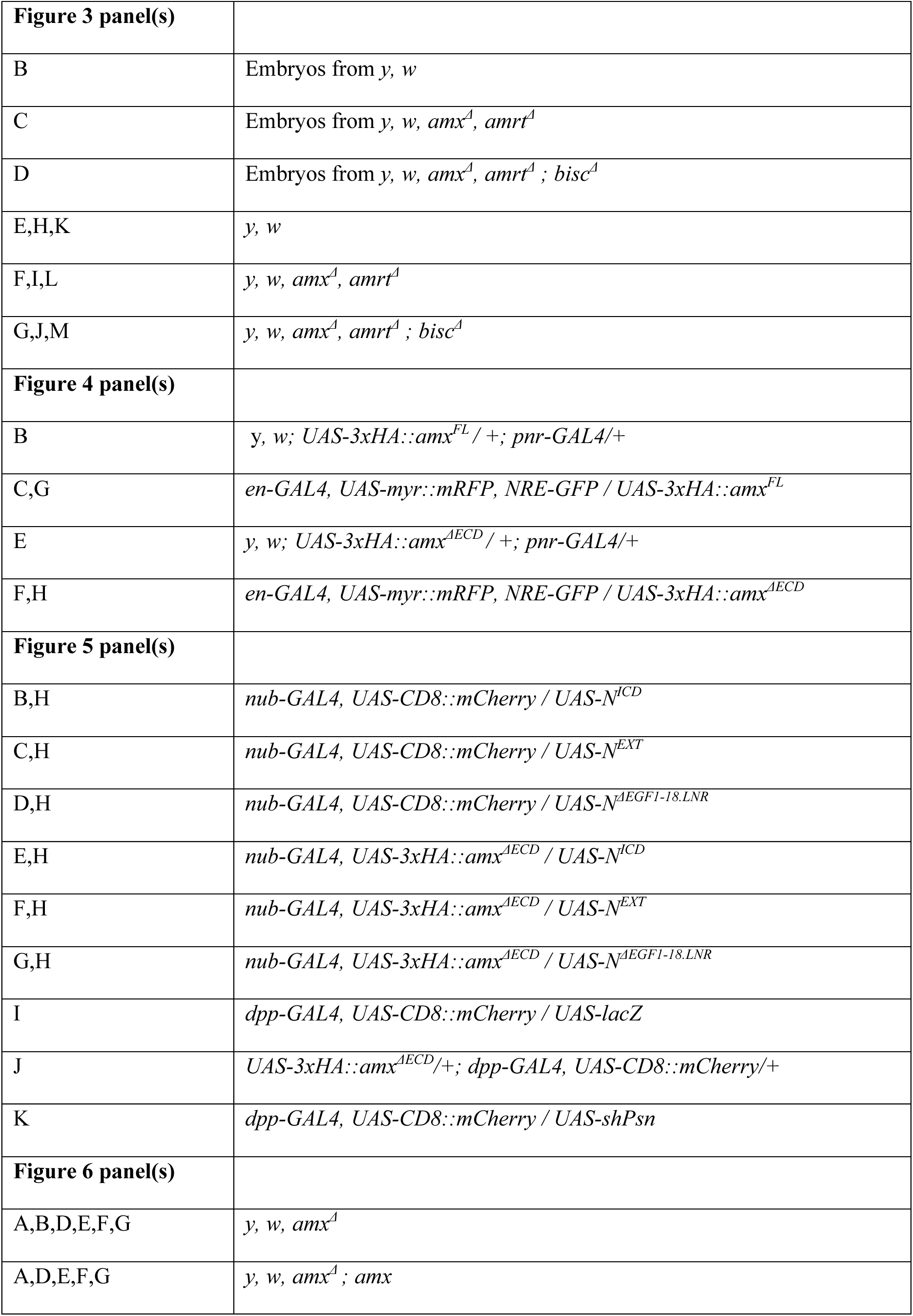
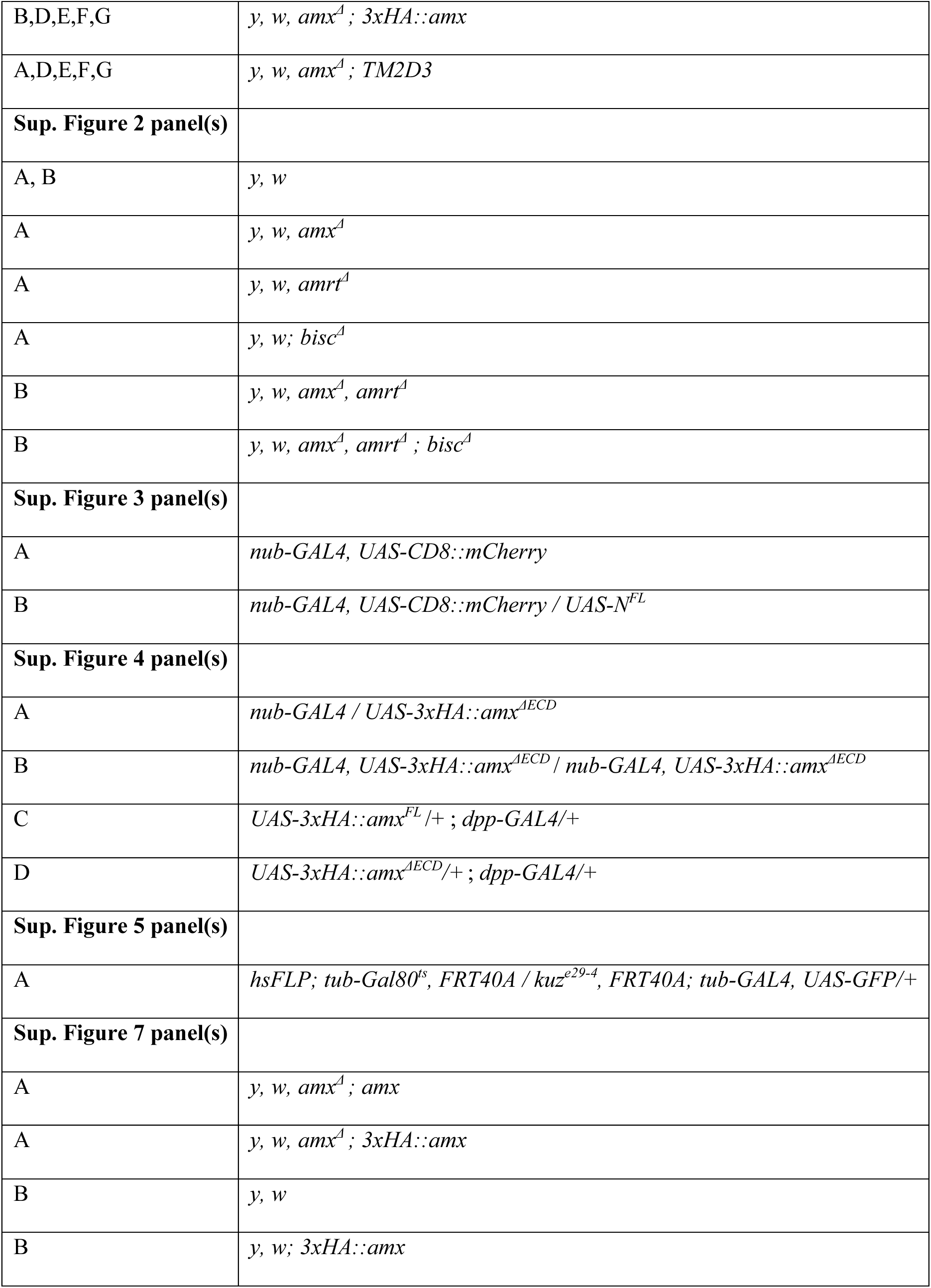

### Generation of N’-3xHA tagged full-length and truncated *amx* overexpression transgenes

The cDNA templates used to generate the UAS constructs for 3xHA epitope tagged full-length and truncated Amx were synthesized by Genewiz®. The *amx* open reading frame was based on the coding sequence of the genomic rescue construct we reported in (Jakobsdottir et al., 2016). A 3xHA epitope sequence (N’-YPYDVPDYAGYPYDVPDYAGSYPYDVPDYA-C’) was inserted after the signal sequence (SS) as predicted by SignalP 4.1 (http://www.cbs.dtu.dk/services/SignalP-4.1) and Phobius (http://phobius.sbc.su.se). A 968 bp fragment corresponding to 3xHA tagged full-length Amx was synthesized with NotI and XbaI restriction sites on the 5’ and 3’ end respectively. Furthermore, a 464 bp fragment corresponding to 3xHA::amx^ΔECD^ was designed based on the full-length 3xHA::Amx construct, but with the majority of the endogenous Amx N’-end removed leaving the SS, epitope tag, and TM2 domains intact. We subcloned the synthesized fragments into pUASTattB (Bischof et al., 2007) via restriction enzyme cloning utilizing the NotI and XbaI cut sites. We then injected the final plasmids into VK37 (*PBac{y[+]-attP}VK00037*) attP docking site (Venken et al., 2006) and injected animals were crossed to *y w* and screened for the presence of the *white*^+^ marker encoded in the pUASTattB backbone in a *white^-^* background. Positive animals were then crossed to *SM6a* to create a balanced stock. One line each was chosen for this study.

### Generation of a N’-tagged *amx* genomic transgene

The 3xHA::amx genomic rescue transgene was designed and constructed based on the non-tagged genomic rescue construct we reported in (Jakobsdottir et al., 2016). A ∼3.3kb region including the full *amx* gene has the ability to fully rescue the maternal effect neurogenic phenotype of *amx^1^* (Jakobsdottir et al., 2016) and *amx^Δ^* (this study). We inserted a 3xHA sequence into the pattB-amx genomic rescue construct described in (Jakobsdottir et al., 2016) via NEBuilder® HiFi DNA Assembly. 3xHA was inserted after the predicted signal sequence (N’-ATMRLQRQCIVVNMRSAIVLIMIFVLTGIRNSET-C’) to tag Amx at its N’. pUASTattB-3xHA::amx^FL^ described above was used as a template to amplify and add appropriate homology arms to the SS-3xHA::Amx DNA sequence with the primers 5’- CCCCGCTCTATCTGACCAAAGCCACCATGAGGCTCCAACGAC-3’ and 5’- AAAACTAAACTAAGAACGGACTACTATATGTAAAGTGAGCCATCCGC-3’ using Q5® High-Fidelity 2X Master Mix (M0492S, NEB). The section of pattB-amx plasmid containing the *amx* regulatory elements was linearized by PCR using Q5 polymerase and primers 5’- CGTTGGAGCCTCATGGTGGCTTTGGTCAGATAGAGCG-3’ and 5’- GCTCACTTTACATATAGTAGTCCGTTCTTAGTTTAGTTTTACAGGGGT-3’, which add appropriate homology arms to allow for assembly with the 3xHA::Amx fragment. The construct was assembled following the protocol described by NEBuilder® HiFi DNA Assembly Master Mix (NEB Catalog #E2621S) and confirmed by Sanger sequencing. The validated construct was injected into embryos expressing ϕC31 integrase with a 2^nd^ chromosome *attP* docking site (VK37) (Venken et al., 2006). Transgenic flies were isolated based on eye color (*w^+^* encoded by the *mini-white* gene in pattB vector) and balanced over *SM6a*.

### Generation of the *amrt^Δ^* mutant

The *amrt (CG11103)* null allele was created based on a method we described in (Li-Kroeger et al., 2018). We selected gRNA (guide RNA) target sites using CRISPR Optimal Target Finder (http://targetfinder.flycrispr.neuro.brown.edu/). sgRNAs (single gRNA) expressing plasmid was generated using the pCFD3-dU6:3 gRNA vector (Plasmid ID: #49410, Addgene) as described in (Port et al., 2014). Oligo DNA to generate the upstream sgRNA plasmid were 5’-GTCGCGCTGCGTGCCTGTATCGCT-3’ and 5’-AAACAGCGATACAGGCACGCAGCG-3’. Oligo DNA to generate the downstream sgRNA plasmid were 5’-GTCGTCCTCGGGCAGCAAATTGT-C’ and 5’- AAACACAATTTGCTGCCCGAGGA-3’. Donor plasmid containing the *yellow^body^*^+^ marker (*y^body^*^+^) to be integrated into the *amrt* locus via HDR was generated through NEBuilder® HiFi DNA Assembly. Primers amplifying and adding homology arms to a pBH vector (Housden and Perrimon, 2016) backbone were 5’- GATGCTGTTAGACTAACGGTGTATATCTAGAGCCGTCCCGTCAAG-3’ and 5’-CCGCAAGCAATGGCCAAACTGGGTCCTCGAGTCGACGTTG-3’. Primers for amplifying the *ybody^+^* insert and adding homology arms from P{ybody+} plasmid (Li-Kroeger et al., 2018) were 5’-CAGACAACACGATGCCCAAGCGGATCGCTTGATGTTGTTTTGTTTTG-3’ and 5’-TGCCGTCCTCGGGCAGCAAATGGAAGGAACCTGCAGGTCAACG-3’. All plasmids were verified by Sanger sequencing. *y, w, iso#6(X); attP2{nos-Cas9}* (Li-Kroeger et al., 2018) embryos were injected with 25 ng/ul concentration of each sgRNA plasmid mixed with 150 ng/ul of the *y^body^*^+^ donor plasmid. Resulting adults were crossed to *y w* animals and offspring screened for the presence of the *y^body+^* marker (dark body color instead of yellow body). Positive animals were crossed to the *FM7c* balancer to establish the lines and were molecularly genotyped (see below).

### Generation of the *bisc^Δ^* mutant

The *bisc* (*CG10795*) null allele was generated using the same strategy discussed above that was used to generate the *amrt* null allele. Oligo DNAs to generate the upstream sgRNA plasmid were 5’- GTCGTATGAGGGACCATGTACAT-3’ and 5’-AAACATGTACATGGTCCCTCATA-3’. Oligo DNA to generate the downstream sgRNA plasmid were 5’-GTCGAGGCGGTGGTGGTGTTCTGT-3’ and 5’-AAACACAGAACACCACCACCGCCT-3’. Primers amplifying and adding homology arms to a pBH vector backbone were 5’-GTAGACACACGGCATAGATGGTATATCTAGAGCCGTCCCGTC-3’ and 5’-GAAGAAGTTGACAATGTGTTGGGTCCTCGAGTCGACGTTG-3’. Primers for amplifying the *ybody^+^* insert and adding homology arms from p{ybody+} plasmid were 5’-AATTGTATGAGGGACCATGTAGGATCGCTTGATGTTGTTTTG-3’ and 5’-GTTGACCTGCAGGTTCCTGTTGGCGGGAGCTCTTTCTC-3’. All plasmids were verified by Sanger sequencing. *y, w; iso#2(2); attP2{nos-Cas9}* embryos were injected with 25 ng/ul concentration of each sgRNA plasmid mixed with 150 ng/ul of the *y^body+^* donor plasmid. Resulting adults were crossed to *y, w* animals and offspring screened for the presence of the *y^body^*^+^ marker. Positive animals were crossed to the *SM6a* balancer to establish the lines and were molecularly genotyped (see below).

### RT-PCR verification of mRNA expression in null mutant flies for *TM2D* genes

The presence or absence of mRNA corresponding to *TM2D* genes were determined using RT-PCR (Reverse Transcription-Polymerase Chain Reaction). Whole body RNA was isolated from adult animals through standard TRIzol/chloroform RNA extraction protocol. We prepared cDNA with iScript™ Reverse Transcription Supermix, (#1708840, BioRad). PCR was done using Q5 polymerase (M0492S, NEB). Primers used to detect the presence of *amx* cDNA were 5’-TCCCCGCTCTATCTGACCAA-3’ and 5’- GCTCTGTTGCCACATTTCCG-3’. Primers to detect the presence of *amrt* cDNA were 5’- CTACGGACTACTGGCGTTCC-3’ and 5’-CCCGTTTGACCGAGACAGAA-3’. Primers to detect the presence of *bisc* cDNA were 5’-CCCCGCGAACTGCAATAAAC-3’ and 5’- CACAACCTGCAGGGCTATCA-3’. Primers targeting *TM2D* cDNA were annealed at 68⁰ C, extended for 30s at 72⁰ C for 30 cycles. Primers for control gene *rp49* were 5’-TCTGCATGAGCAGGACCTC-3’ and 5’-CGGTTACGGATCGAACAAG-3’ (Li et al., 2014); annealed at 64⁰ C and extended 30s for 30 cycles.

### Generation of an *amrt* genomic rescue construct

Genomic DNA was isolated from *y, w, iso#6; +/+; attP2{nos-Cas9}* animals using PureLink Genomic DNA Mini Kit (Cat no. K1820-01, Invitrogen). Genomic region fully containing the *amrt* (*CG11103*) locus and neighboring sequences was amplified by PCR using the primers 5’- TATATACTCGAGcgcgaaactttcgatttcc-3’ and 5’-TATATAGAATTCatcgaatgtagagatgggc-3’ (small letters indicate annealing region) which added EcoRI and XhoI restriction sites for follow-up cloning into the pattB vector (Bischof et al., 2007). The pattB-amrt plasmid was injected into VK37 (Venken et al., 2006) docking site on the second chromosome. Eclosed animals were crossed to *y, w* and screened for the *w*^+^ marker in the subsequent generation. Positive animals were then crossed to *SM6a* to create a balanced stock.

### Generation of *biscotti::GFP* fosmid transgenic line

A ∼40kb genomic fosmid construct in which *bisc* is C’-tagged with GFP and other epitopes (Sarov et al., 2016) (*bisc::GFP*, FlyFos021003, *Drosophila* TransgeneOme Resource ID: CBGtg9060C1139D) was obtained from Source BioScience. Bacterial colonies were provided to GenetiVision Corp. for DNA preparation, injection into VK33 (*PBac{y[+]-attP}VK00033*) on the third chromosome (Venken et al., 2006), selection, and balancing of fly lines using the *TM3, Sb* balancer. Three independent lines were generated and all three behaved in a similar manner. One line was chosen for the experiments performed in this study.

### Creation of Notch overexpression transgenic lines (*UAS-Notch*)

All transgenic constructs were generated by Gateway® (Thermo Fisher Scientific) cloning into the pUASg-HA.attB plasmid (Bischof et al., 2007). First, we generated Gateway compatible plasmid that contains the full-length Notch open reading frame. We subcloned the full-length *Notch (N^FL^*) open reading frame into the pDONR223 plasmid based on a cDNA clone provided by Dr. Spyros Artavanis-Tsakonas (Wharton et al., 1985), which was mediated by a Gateway reaction using BP clonase II (Thermo Fisher Scientific, #11789100) following a PCR reaction and addition of attB sites to the amplicon (Harnish et al., 2019; Marcogliese et al., 2018). Truncated *Notch* constructs were generated by Q5 site-directed mutagenesis (NEB) with the following primers:

*N^EXT^* : 5’-GCGGCCAAACATCAGCTG-3’ and 5’-GTGCATTTTGTTAATCCAAAAACAAATCC-3’.

*N^ICD^* : 5’-GTCTTGAGTACGCAAAGAAAG-3’ and 5’-CATGGTGAAGCCTGCTTT-3’.

*N^ΔEGF1-18.LNR^* (two-step mutagenesis): 5’-CTGAGCGATGTGGACGAGTGCGCATCGAAT-3’ and 5’-CAACGCGGTATCAGTTCC-3’ followed by mutagenesis with 5’-AACAAGACCCAGTCACCG-3’ and 5’-CATGGCACGTTGTTGCTC-3’.

All constructs were fully sequenced (Sanger), and cloned into pUASg-HA.attB via LR clonase II (Thermo Fisher Scientific, #11791020). All expression constructs were inserted into VK37 integration site (Venken et al., 2006). Two transgenic lines were established for each and one line each was used for this study.

### Embryo collection, staining and imaging

Embryo collection, staining and imaging was performed as previously described (Jakobsdottir et al., 2016). In brief, virgin females that are homozygous for each or all *TM2D* gene mutations, with or without genomic rescue constructs, were crossed to males flies of the same genotype or *Canton-S* males, and allowed to mate in a vial for 24 hrs. Flies were then transferred to a bottle with a grape juice plate supplemented with active yeast and allowed to lay eggs overnight. Embryos were then gently collected using a paint brush and their chorions were removed by 1.5 minute incubation in 66% bleach. Dechorionated embryos were then washed with water and fixed in 4% paraformaldehyde/PBS(phosphate buffered saline)/n-heptane solution for 30 minutes at room temperature. Fixed embryos were washed and stored in 100% methanol at -20°C until use. Fixed embryos were rehydrated and rinsed with 0.03% Triton-X in PBS (PBST). A primary antibody to label neuronal nuclei [anti-Elav, rat monoclonal (7E8A10) (O’Neill et al., 1994), 1:200, Developmental Studies Hybridoma Bank (DSHB)] was applied in a solution of PBST/5% normal donkey serum (NDS)/0.1% NaN_3_ overnight at 4°C. Embryos were further washed with PBST upon removal of the primary antibody, and a secondary antibody (donkey anti-rat-Alexa488, 1:500; Jackson ImmunoResearch #712-545-153) was applied for 1 hour at room temperature. Stained embryos were washed in PBST and mounted onto glass slides using Vectashield^®^ with DAPI (4’,6-diamidino-2-phenylindole, Vector labs). Embryos were imaged using Ti2E Spinning Disc confocal microscope (Nikon) and images analyzed using NIS software (Nikon).

### Egg hatching assay

Egg hatching assay was performed as previously described (Jakobsdottir et al., 2016). Embryos were collected and dechorionated as described above. Dechorionated embryos were suspended in PBS and placed in 12-well cell culture dishes. The dishes were then placed in a 25°C incubator for 24 hours. The ratio of hatched to unhatched embryos after the 24 hour period was recorded for each genotype and used to calculate hatching rate (%). This was repeated at least 3 times and statistical analysis and graph generation was performed using GraphPad Prism 9.0 software. We performed one-way ANOVA followed by Dunnett test or t-test. **** = p-value ≤0.0001.

### Imaging of adult flies

Heads and wings from adult flies were removed from the body using fine dissection scissors and imaged directly using a MZ16 microscope (Leica) with attached Microfire camera (Optronics) using ImagePro Plus 5.0 acquisition software (Media Cybernetics). Extended focus function was used to obtain deep focus images out of Z-stack images. Imaging of the dorsal thoraxes (nota) were described in (Yamamoto et al., 2012) with slight modifications. In brief, legs, head, and abdomen were removed from thoraxes with fine dissection scissors. Then, the dissected thoraxes were then placed in 10% KOH at 95°C. for 10 min to dissolve soft tissue. Thoraxes were then further trimmed with scissors prior to imaging and mounted on a glass slide with spacers using 75% glycerol/25% ethanol solution. Photos were taken using microscope system and imaging software as above.

### Immunostaining and imaging of *Drosophila* wing imaginal discs and ovaries

Wing discs from wandering larvae and ovaries from females were dissected in 1x PBS and fixed for 30 minutes in 4% paraformaldehyde in PBS. Tissues were then washed with 0.2% PBST. Primary antibodies [mouse anti-Notch intracellular domain (NICD) (1:50; DSHB, C17.9C6), mouse anti-Cut (1:100; DSHB, 2B10), rat anti-HA (1:100, Sigma-Aldrich, 11867423001)] were applied in 0.2% PBST with 5% NDS/0.1% NaN_3_ overnight at 4°C. Tissue was then washed with 0.2% PBST 3 times, 15 minutes and secondary antibodies/stains [donkey anti-rat IgG-Cy3 (1:500; Jackson ImmunoResearch #712-165-153) donkey anti-mouse IgG-Alexa-647 (1:500; Jackson ImmunoResearch #715-605-151), donkey anti-mouse Alexa-488 (1:500; Jackson ImmunoResearch #715-545-151), and Alexa-488 Phalloidin (1:1000; ThermoFisher A12379)] were applied in 0.2% PBST/NDS for 2 hours at room temperature. Tissues were washed with 0.2% PBST and mounted in Vectashield® with DAPI (Vector labs). Images were taken with LSM 710 Confocal Microscope (Zeiss).

### Notch epistasis assay

*UAS-Notch* lines were crossed to either *nub-GAL4, UAS-CD8::mCherry* (control) or *nub-GAL4, UAS-amx^ΔECD^* flies. Wing discs from 3^rd^ instar larvae were dissected out, fixed washed and stained for Cut as described earlier. Cut intensity within the wing pouch was quantified using ImageJ. Graph generation and statistical analysis was performed with GraphPad Prism software version 9.0. One way t-test was used to compare experiments. *= p≤0.05. ****= p≤0.0001

### MARCM analysis

The following fly lines were used for MARCM (Lee and Luo, 1999) analysis:

*hsFLP; tub-Gal80^ts^, FRT40A/CyO; tub-GAL4, UAS-GFP/TM6b, Tb* (Yang and Deng, 2018)
*kuz^e29-4^, FRT40A/CyO* (Rooke et al., 1996)

Flies were crossed and maintained at 25°C. First-instar larvae (36-48 hours after egg laying) were heat shocked for 40 minutes twice a day at 37°C. Wing discs were dissected in 1×PBS from wandering larvae (108-120 hours after egg laying). Discs were then fixed in 4% paraformaldehyde in PBS for 20 minutes at room temperature. Fixed discs were washed with 0.2% PBST. A mouse anti-Notch primary antibody (1:40; DSHB, C17.9C6, raised against the intracellular domain) was applied in 0.2% PBST with 5% NDS/0.1% NaN_3_ overnight at 4°C. Discs were further washed with 0.2% PBST after primary antibody staining. A donkey anti-mouse-Cy3 secondary antibody (1:500; Jackson ImmunoResearch #715-165-151) in 0.2% PBST was applied for 2 hours at room temperature. Stained discs were washed with 0.2% PBST and mounted with Vectashield® with DAPI (Vector labs). Fluorescence Images were taken by LSM 710 Confocal Microscope (Zeiss).

### Western blot of 3xHA::Amx

To determine whether 3xHA::Amx is expressed in adult brains or ovaries, we performed western blot. Adult brains from *y, w; VK37{pattB-3xHA::amx}* flies were dissected out as described in (Tito et al., 2016). For ovaries, we mated the *y, w; VK37{pattB-3xHA::amx}* females flies to male flies of the same genotype while supplying plenty of active yeast to stimulate oogenesis for two days prior to protein isolation. Ovaries were then dissected out from the abdomen in cold (4°C) PBS. Dissected brains and ovaries were rinsed with cold PBS and placed immediately in cold 8M urea lysis buffer (8M urea, 10% glycerol, 0.5% SDS, 5% β-mercaptoethanol) with Halt™ Protease Inhibitor Cocktail 100X (Thermo Scientific, #78430) added before lysis. We homogenized the brains or ovaries were via pestle in 15 uL of 8M urea lysis buffer. Homogenate was incubated for 30 minutes on ice, and 2x Laemmli Sample Buffer (Bio-Rad, #1610737) was added prior to gel loading. Best results were obtained when avoiding heating/boiling protein sample. Homogenate was loaded directly into Mini-PROTEAN® TGX™ 4-20% Precast Gels (Bio-Rad, #4561094). SDS-PAGE (Sodium Dodecyl Sulphate–PolyAcrylamide Gel Electrophoresis) was run for 30 minutes at 120V in Tris/Glycine/SDS buffer (BioRad, #1610732). Protein was then transferred onto PVDF membrane using Bio-Rad TransBlot Turbo system using the low molecular weight protocol. The membrane was blocked with 5% skim milk in 0.5% Tween-20/Tris-Buffered Saline (TBST) for 1 hour at room temperature and then washed 3 times for 5 minutes with TBST. A primary antibody (rat anti-HA, 1:1,000, Sigma, 3F10) was diluted in 5% fetal bovine serum/TBST and the membrane was incubated overnight at 4°C. The membrane was again washed with TBST and a secondary antibody (donkey anti-rat HRP, 1:5,000, Jackson ImmunoResearch, #712-035-150) was applied in 5% milk/TBST for 2 hours at room temperature. The membrane was washed again and imaged using SuperSignal™ West Femto Maximum Sensitivity Substrate (Thermo Scientific, 34096) and ChemiDoc imaging system (Bio-Rad) using default settings.

### Longevity assay of *amx^Δ^* flies

To determine whether loss of *amx* causes lifespan defects, we compared the longevity of male flies that lack *amx* (*y, w, amx^Δ^*) to flies in which *amx* function has been rescued with genomic constructs that express wild-type Amx (*y, w, amx^Δ^*; *VK37{pattB-amx}*), N’-3xHA tagged Amx (*y, w, amx^Δ^; VK37{pattB-3xHA::amx}*) or human TM2D3 expressed under the control of fly *amx* regulatory elements (*y, w, amx^Δ^; VK37{pattB-TM2D3}*). Flies were reared and collected as described in (Linford et al., 2013). Ten animals were housed together in a single vial and flies were flipped to a new vial with fresh food every 2-3 days. Vials were kept in a 25°C incubator with a 12 hr light/dark cycle. Dead flies were recorded after every vial flip. Generation of the Kaplan-Meier curve and statistical analysis was performed with GraphPad Prism 9.0. We applied log-rank test (Mantel-Cox), ****= p≤0.0001.

### Electrophysiological recordings of the giant fiber system

Electrophysiological recordings of the giant fiber system were performed with a protocol modified from (Tanouye and Wyman, 1980). Flies were first anesthetized on ice then transferred to a petri dish filled with soft dental wax; wings and legs were mounted in wax, ventral side down, using forceps. Five electrolytically sharpened tungsten electrodes were used: two for stimulating the giant fiber, one as a reference electrode, and two for recording from the TTM and DLM, respectively. To activate the giant fiber, two sharp tungsten electrodes were inserted into each eye and voltage stimulation was applied at different frequencies ranging from 0.5 Hz to 100 Hz. DLM and TTM responses were measured through the two electrodes implanted in the DLM and TTM.

For each adult fly, prior to applying high frequency stimulation on the giant fiber, low frequency stimulations at 0.5 Hz were applied after placing the two recording electrodes in TTM and DLM to ensure that the electrodes are recording from the proper muscles (the latency of responses for TTM: 0.8 ms and for DLM: 1.2 ms (Tanouye and Wyman, 1980). For the actual experiments, high frequency train stimulations of 20 pulses were delivered to the giant fiber at 20, 50 and 100 Hz in random order. Ten times repetitive stimulations were applied for each particular frequency train, interspersed with few minutes rests between two trains of stimuli (for 20 Hz, 1 minute resting; 50 Hz 2 minutes and 100 Hz 3 minutes). 0.5 Hz stimulations were used again after high frequency stimulation to confirm that electrodes were still in the proper muscle. The aforementioned process was considered as one biological sample. Stimuli of the crossing electrodes were fixed at a duration of 10 microseconds at 10–13 V of amplitudes through a stimulus isolation unit (Digitimer Ltd, model DS2A) and the frequency of train stimuli was controlled by LabChart Pro-8 acquisition software (ADInstruments). A microelectrode amplifier (A-M system, Model 1800) was used for all recordings. PowerLab 4/35 (ADInstruments) was used for data acquisition. The probability of responses for one biological sample, under particular frequency of giant fiber stimulation, due to a particular stimulus, was calculated from the proportion of successful responses (out of 10) for both TTM and DLM pathways. The difference of ‘probability of responses’ between control and experimental samples (p-value) for each stimuli were calculated by multiple unpaired t-tests with Holm-Šídák correction for multiple comparisons using Graph Pad Prism 9.0. n.s.= not significant. * = p≤0.05. ** = p≤0.01. *** = p≤0.001. **** = p≤0.0001.

## Supplemental Figures and Figure Legends

**Supplemental Figure 1.**
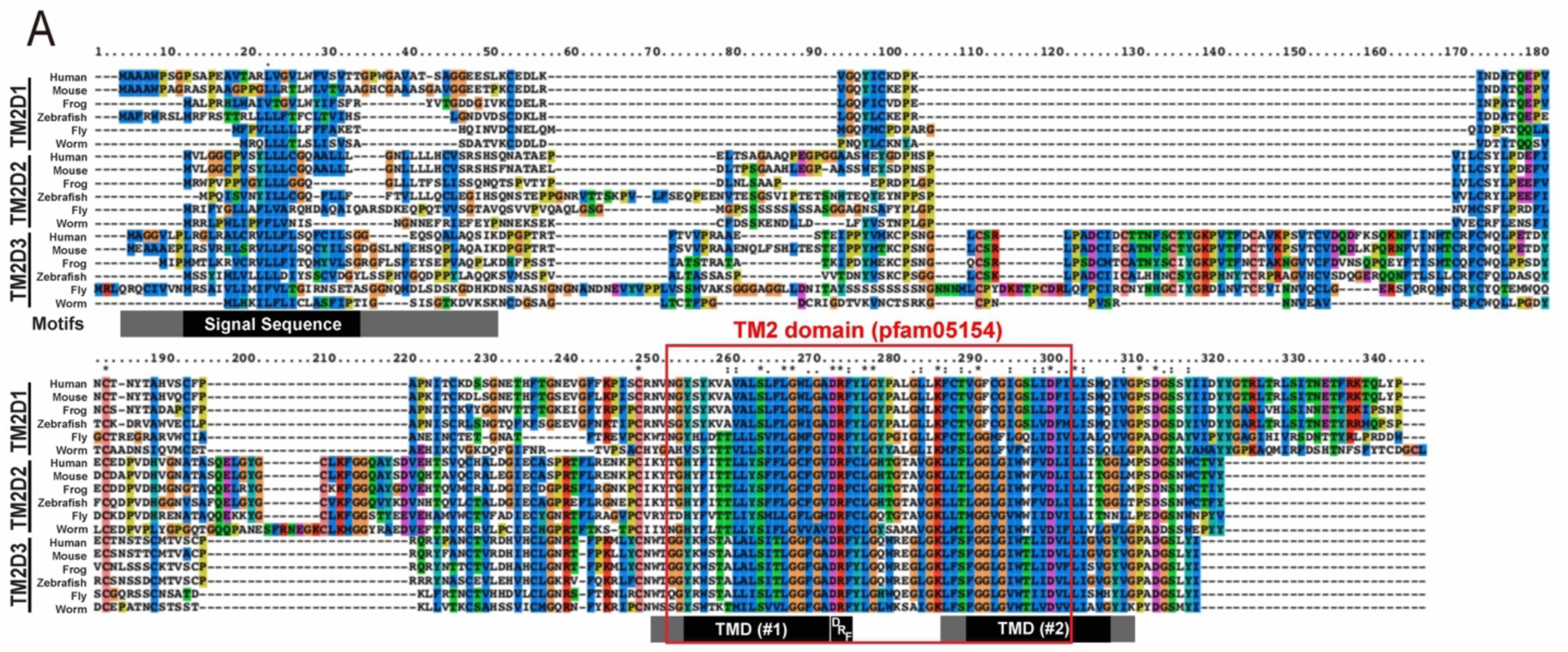
T*M*2D genes are conserved in metazoan species. Protein alignment of Human TM2D proteins across multiple species (human, mouse, frog, zebrafish, fly, worm). TM2 domain (red box, <u>pfam05154</u>) is highly conserved among the proteins and across species. C’ terminus of the proteins are also well conserved across species. Black bars denote the regions that are commonly annotated as Signal Sequence, TMD (Transmembrane domain, #1), DRF or TMD (#2) in all human TM2D1-3 proteins (consensus regions) based on Uniprot (https://www.uniprot.org/). Gray bars show regions that have been annotated as Signal Sequence, TMD (#1) or TMD (#2) in one or two human TM2D1-3 proteins.

**Supplemental Figure 2.**
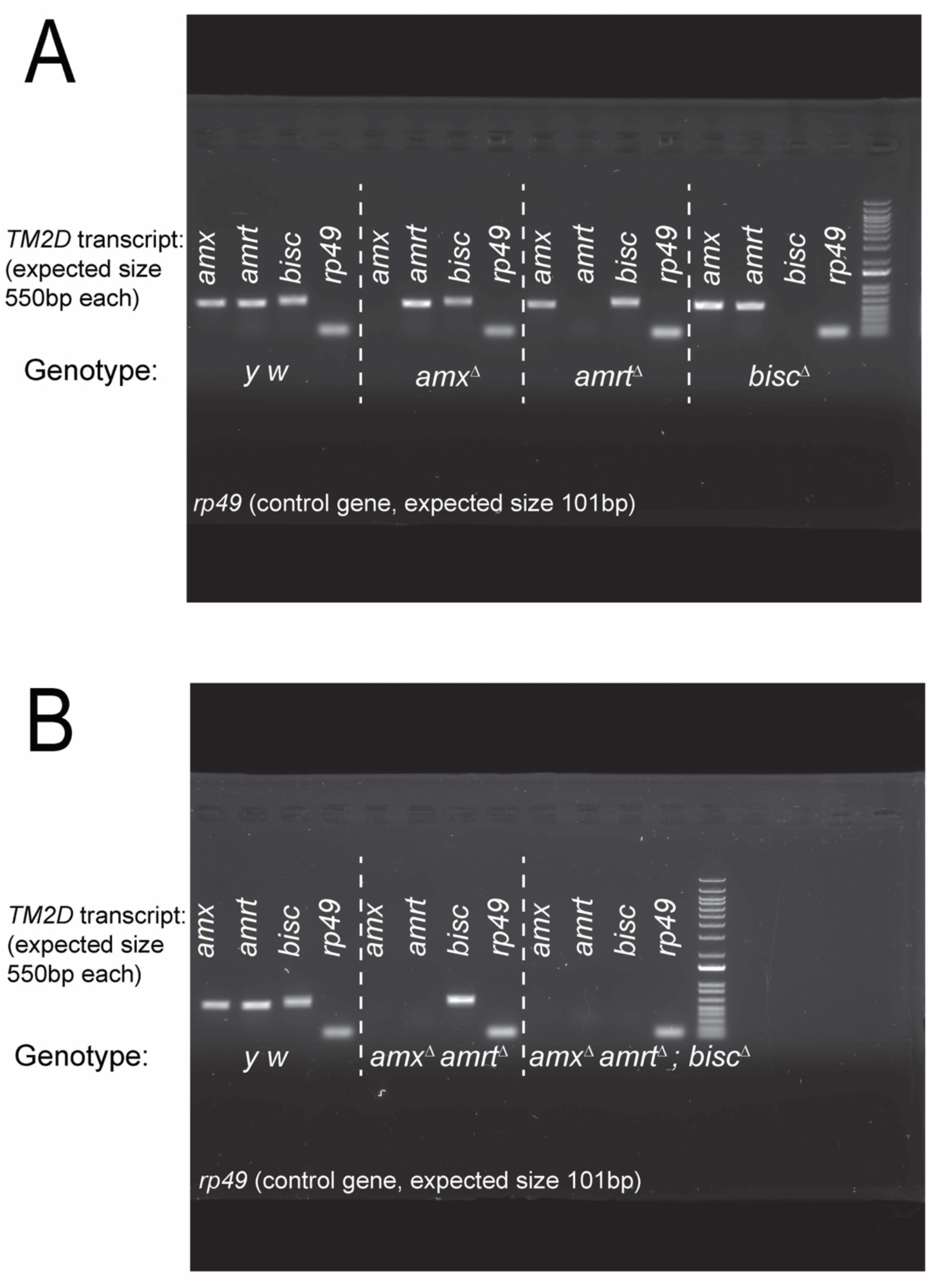
T*M*2D null fly mutants do not express corresponding mRNAs. Reverse transcription followed by PCR (RT-PCR) to verify loss of *TM2D* gene transcripts in mutant fly lines. mRNA was isolated from animals homozygous for their respective alleles. (A) Single mutant lines lack their appropriate gene transcript while other *TM2D* transcripts are unaffected. (B) *amx amrt* double mutants express *bisc*. *TM2D* triple mutants lack all transcripts. *rp49* is a house-keeping gene used as a control for the reverse transcription reaction.

**Supplemental Figure 3.**
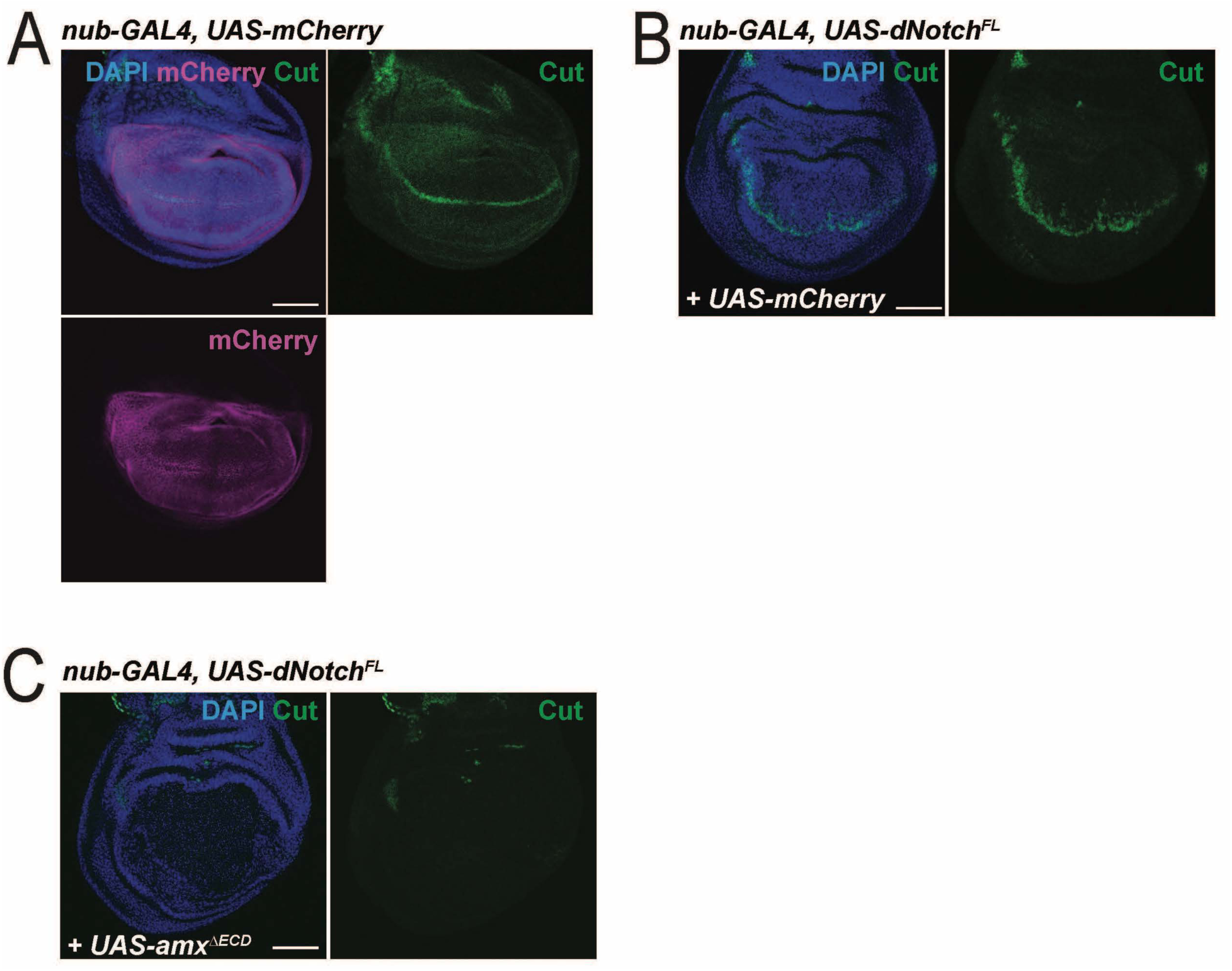
Epistasis experiments between Amx^ΔECD^ and full-length Notch in the wing imaginal disc. (A) A developing wing disc exhibiting normal expression of Notch target Cut (green) within the wing pouch labeled by *UAS-CD8::mCherry* driven by *nub-GAL4* (magenta). (B) Overexpression of full-length Notch in the developing wing pouch via *nub-GAL4* causes a minor upregulation of Cut expression close to the wing margin, likely reflecting the availability of ligands within the wing pouch. (C) Amx^ΔECD^ inhibits the increase of Cut expression induced by Notch as well as abolishing the normal expression levels of Cut, showing it is epistatic to full-length Notch.

**Supplemental Figure 4.**
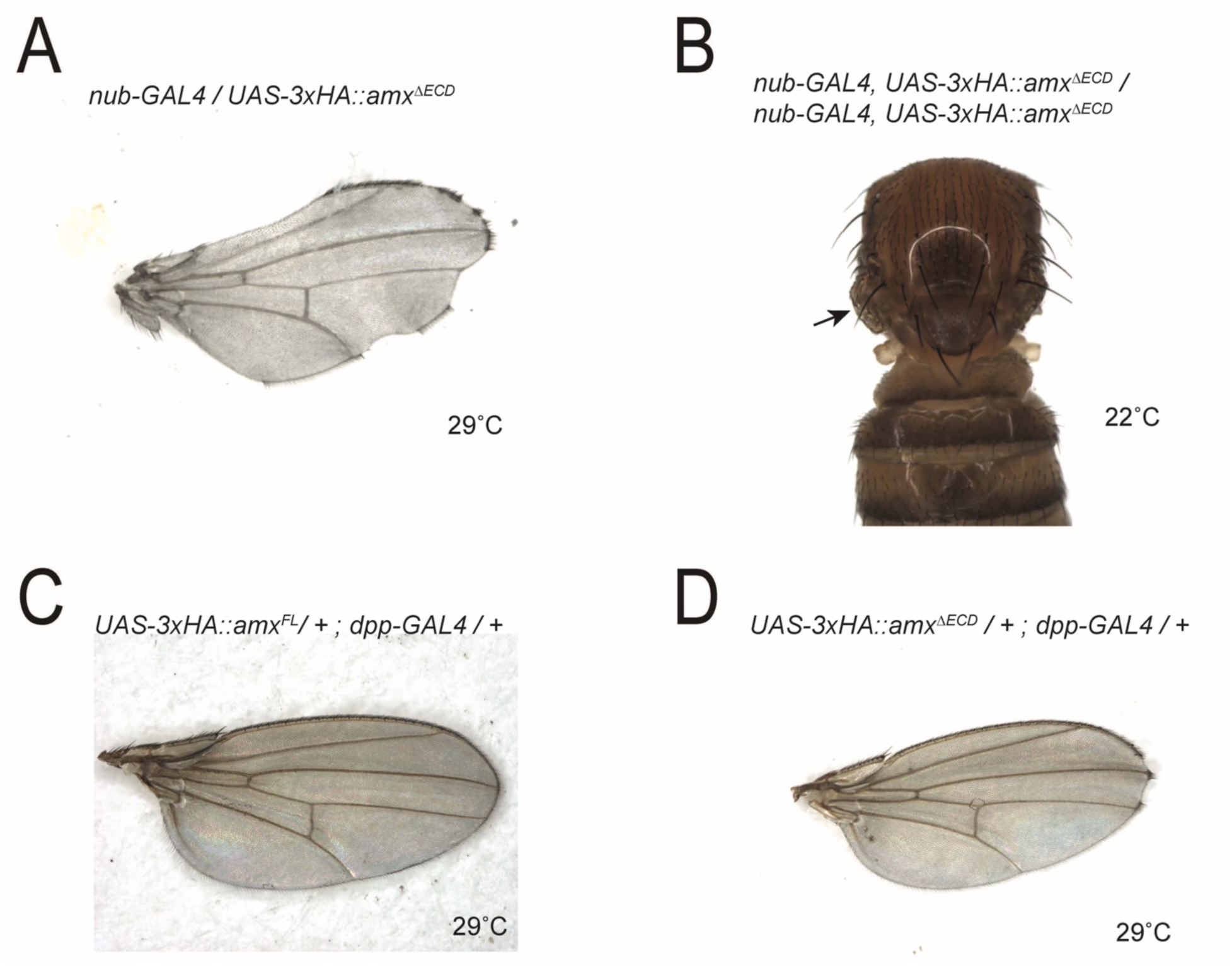
Truncated Amx causes wing notching or wing loss when expressed with multiple wing-expressed GAL4 drivers. (A) Amx^ΔECD^ expressed in the developing wing pouch with *nub-GAL4* causes notching along the wing margin. (B) Homozygous *nub-GAL4, UAS-amx^ΔECD^* recombinant animals show near complete loss of wing (arrow). (C) *dpp-GAL4* (expressed between the third and fourth wing veins) driven expression of Amx^FL^ has no effect on wing morphology. (D) *dpp-GAL4* driven expression of Amx^ΔECD^ causes notching at the wing tip.

**Supplemental Figure 5.**
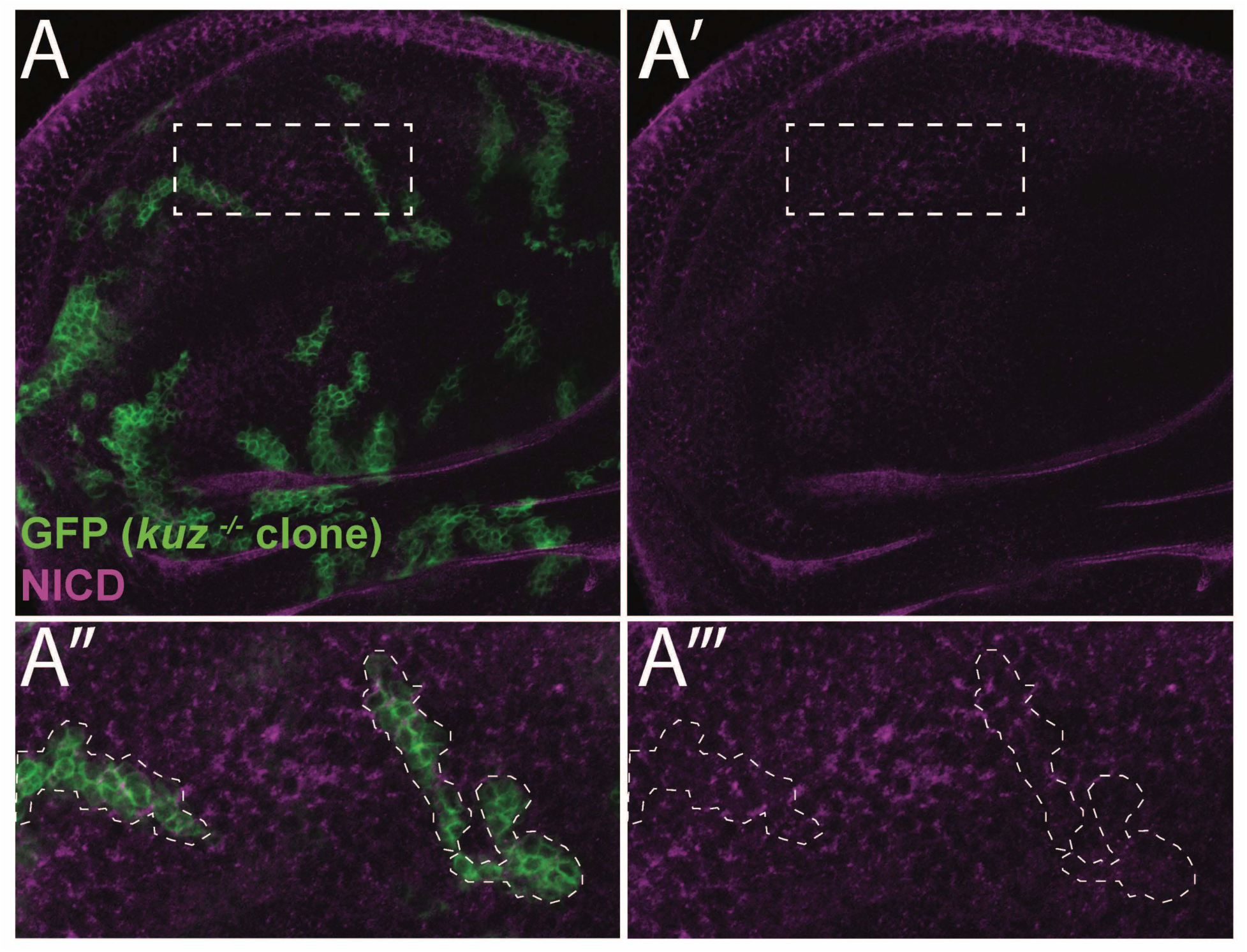
k*u*z (ADAM10) null mutant clones do not show Notch accumulation. (A) *kuz^-/-^* clones (positively marked by GFP, green) were generated by MARCM using a heat shock induced Fippase (*hs-FLP*). The expression level and gross subcellular localization of Notch (magenta) is not altered in *kuz^-/-^* clones compared to control tissue (non-GFP cells). A’’ and A’’’ show the boxed region in A and A’.

**Supplemental Figure 6.**
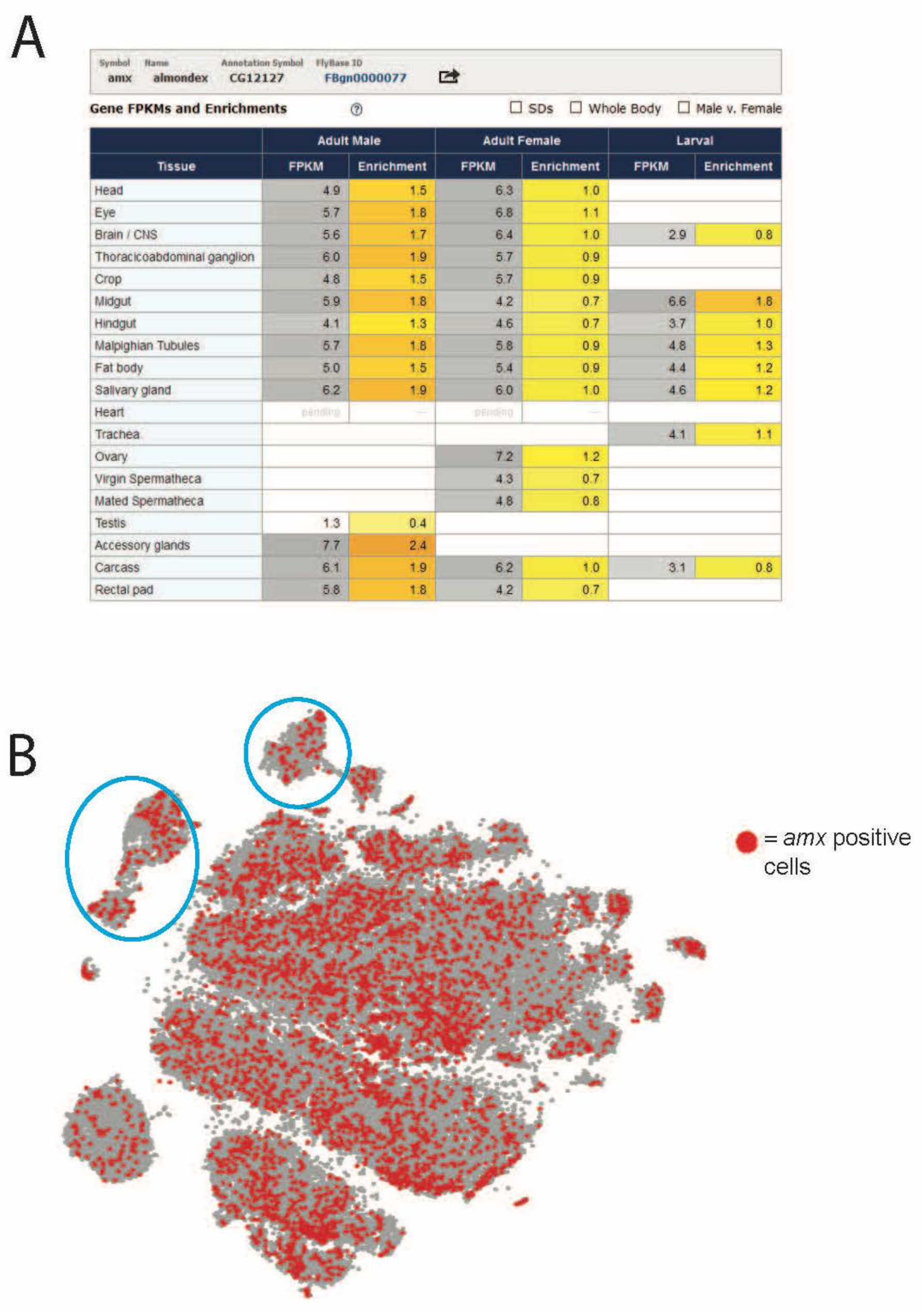
a*m*x mRNA is expressed in nervous system of *Drosophila* according to transcriptomic databases. (A) Summary table for *amx* transcript expression provided by FlyAtlas (http://flyatlas.gla.ac.uk/FlyAtlas2/index.html?search=gene&gene=CG12127&idtype=cgnum#mobileTargetG). *amx* transcript is found in the Brain/CNS of adult flies, as well as other tissues. (B) Single-cell transcript data shows *amx* expressed in many but not all cells in the adult fly brain based on (Davie et al., 2018). Clusters of cells positive for *repo* (glial marker) expression are circled in blue; the remaining cells are largely *elav* (neuronal marker) positive (https://scope.aertslab.org/). Red dots are cells positive for *amx* expression.

**Supplemental Figure 7.**
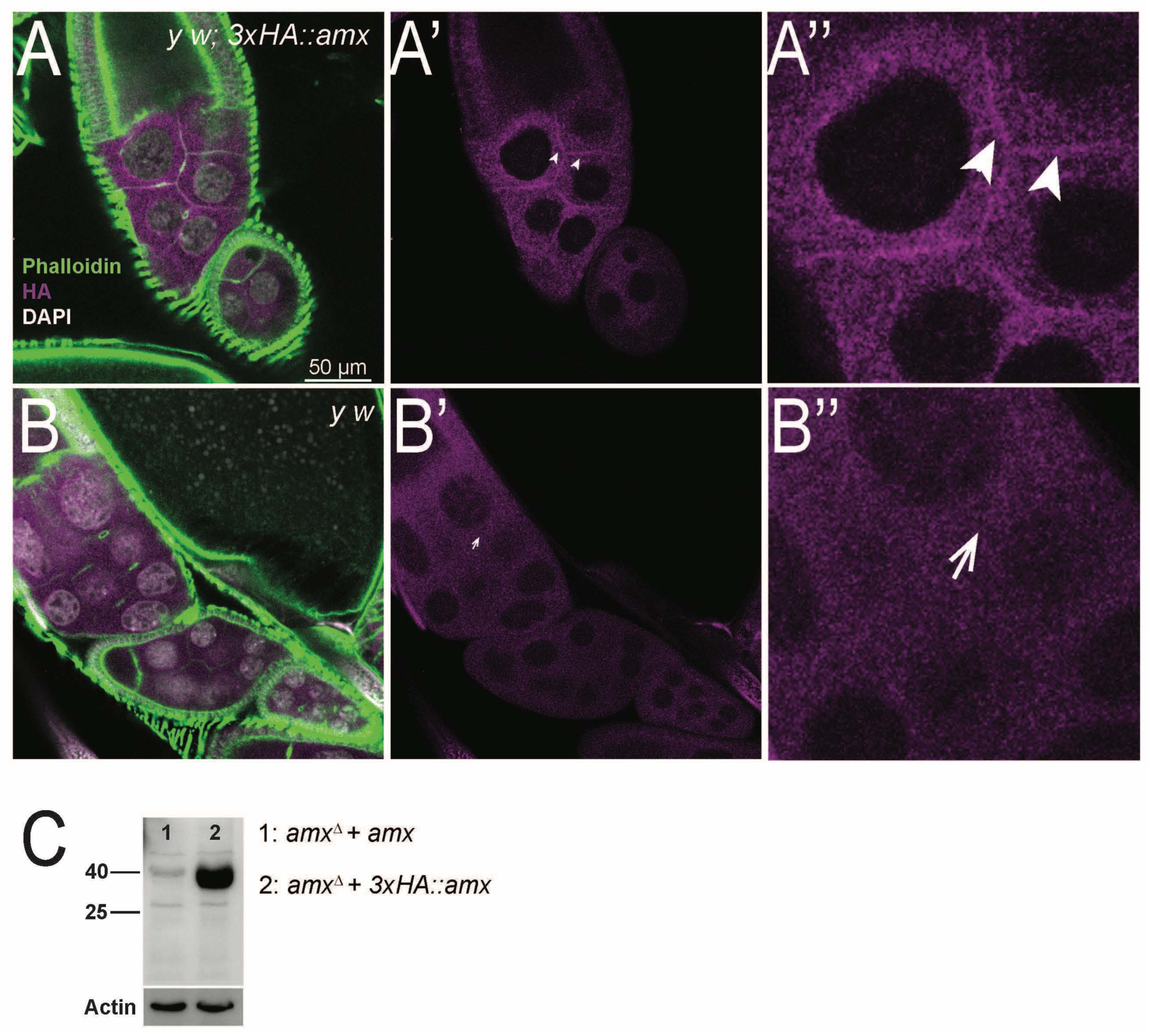
3xHA::Amx is expressed in the *Drosophila* ovary and localizes to the cell membrane as well as intracellular puncta. (A-B) 3xHA::Amx (magenta) localizes to the plasma membrane (marked by Phalloidin, green) separating nurse cells (arrow heads). The signal is relatively low but clearly above background levels of *y w* control (A’’ vs. B’’). The same membranous localization of HA staining is not seen in negative control (arrows). (C) Western blot on ovaries showing positive expression of 3xHA::Amx (lane 2, expected size 35 kDa) in *amx^Δ^* flies compared to untagged Amx control in the same genetic background (lane 1); two ovary pairs were loaded per lane.

**Supplemental Figure 8.**
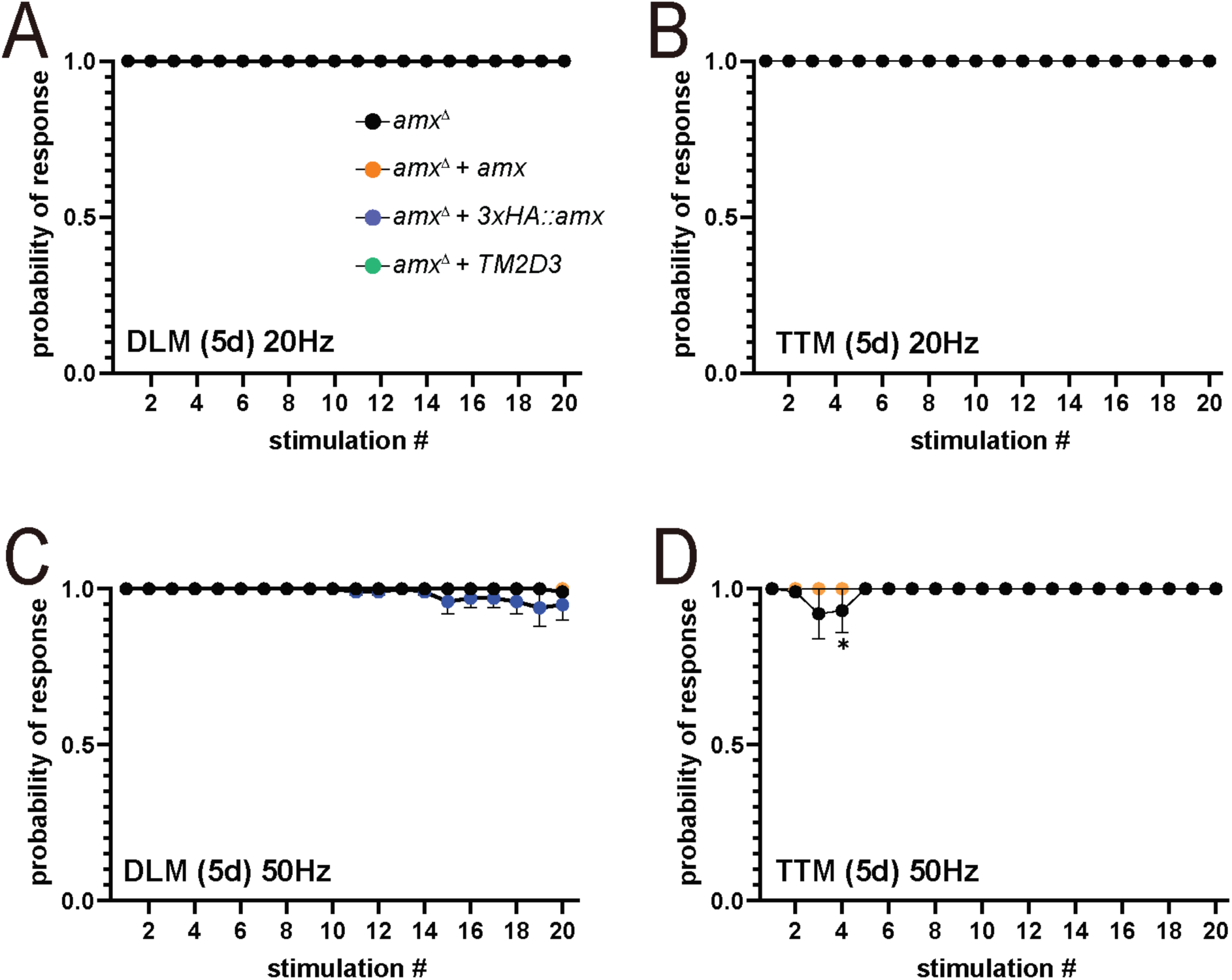
Giant fiber recordings from 5 day post eclosion flies stimulated at 20 and 50 Hz. (A,C). DLM muscles of 5 day old *amx^Δ^* mutants (black) have a response similar to *amx^Δ^ + amx* controls (orange) at stimulation frequencies of 20 and 50 Hz. (B,D) TTM muscles show a small but significant decrease in response probability at 50 Hz but not 20 Hz. *amx^Δ^ + 3xHA::amx* (blue) flies also perform as well as controls (A-D). Multiple unpaired t-tests with Holm-Šídák correction for multiple comparisons. *= p<0.05. Error bars show SEM.

**Supplemental Figure 9.**
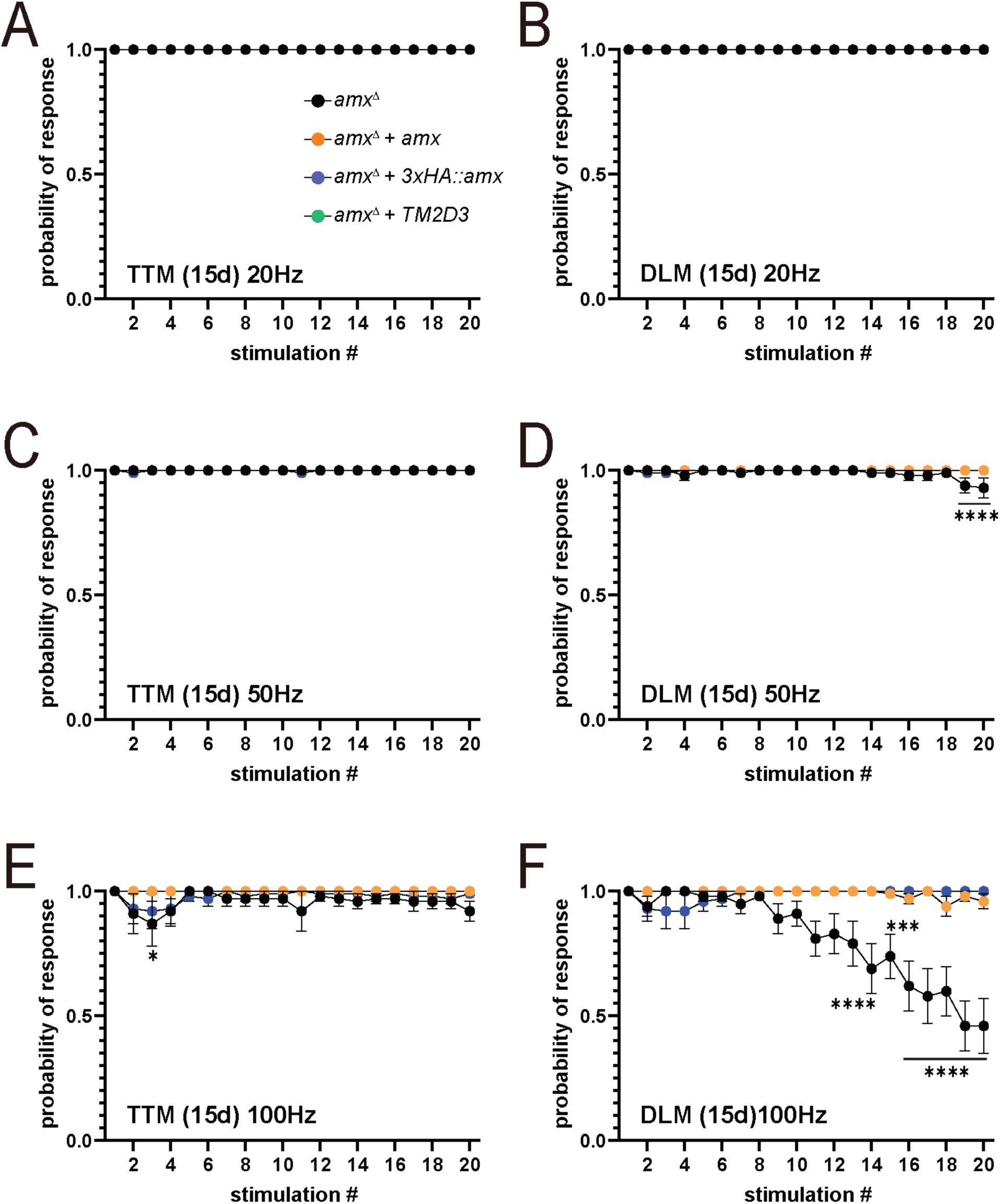
Giant fiber recordings from 15 day post eclosion flies stimulated at 20, 50 and 100 Hz. (A,C,E) TTM failure rate at 20 and 50 Hz. Responses are similar between *amx^Δ^* mutants (black) and *amx^Δ^ + amx* controls (orange) (A, C), with slight but significant failures were observed at 100 Hz (E). (B,D,F) DLM failure rate at 20 and 50 Hz. *amx^Δ^* mutants have a response similar to *amx^Δ^ + amx* controls at 20 Hz (B) but begin to show significant failure to respond at 50 and 100 Hz (D,F). Multiple unpaired t-tests with Holm-Šídák correction for multiple comparisons. ***= p<0.001, ****= p<0.0001. Error bars show SEM.

**Supplemental Figure 10.**
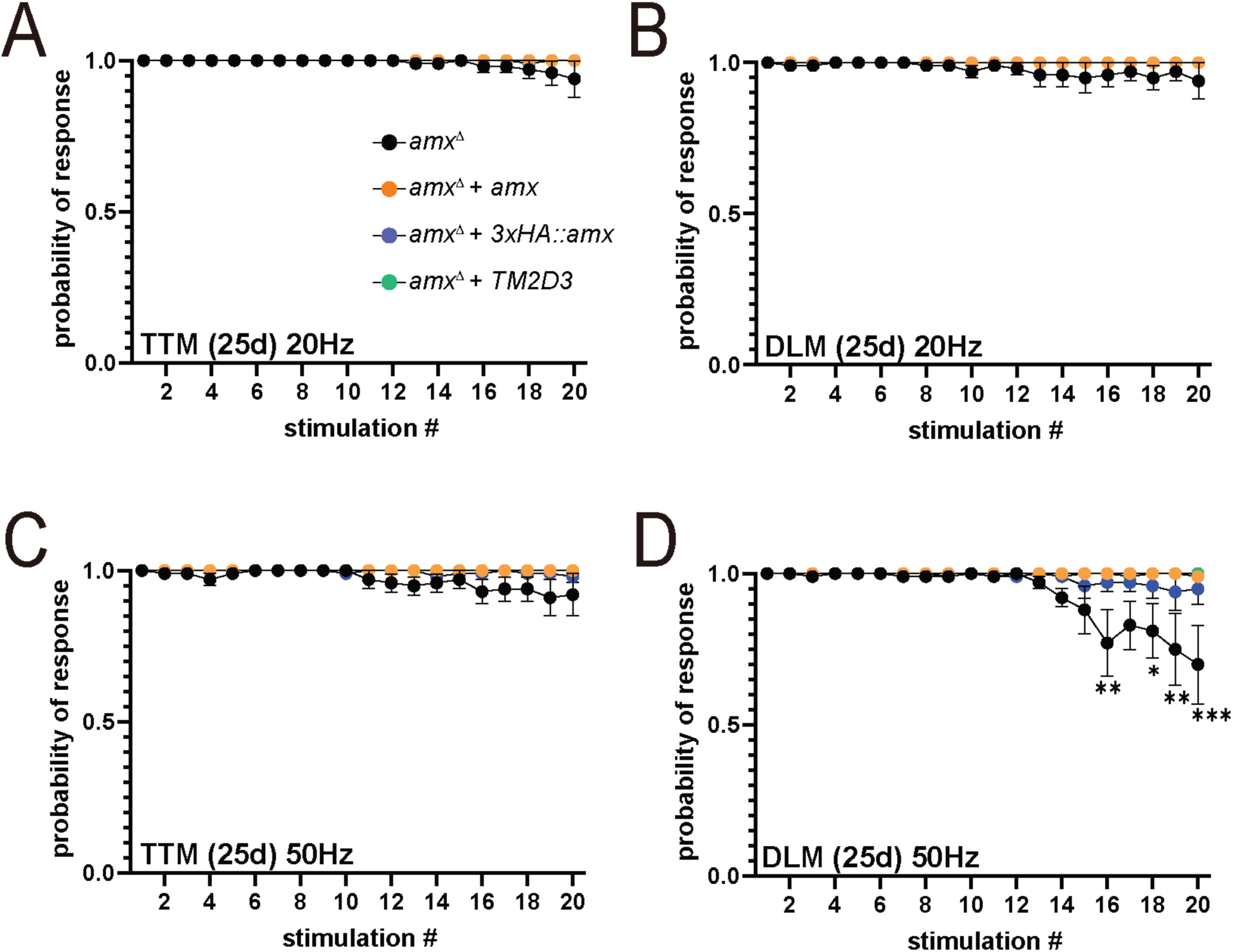
Giant fiber recordings from 25 day post eclosion flies stimulated at 20 and 50 Hz. (A,C) TTM response at 20 and 50 Hz of 25 day old *amx^Δ^* mutants is similar to controls and animals carrying a human *TM2D3* rescue construct. (B,D) At 25d old, *amx^Δ^* mutants perform similarly to controls at 20 Hz (B) but show a significant increase in response failure at 50 Hz (D). Human *TM2D3* rescued flies again perform similarly to controls. Multiple unpaired t-tests with Holm-Šídák correction for multiple comparisons. *= p<0.05. **= p<0.01, ***= p<0.001. Error bars show SEM.

## Key Resources Table

PMID: PubMed ID (https://pubmed.ncbi.nlm.nih.gov), BDSC: Bloomington Drosophila Stock Center ID (https://bdsc.indiana.edu), DSHB: Developmental Studies Hybridoma Bank ID (https://dshb.biology.uiowa.edu)

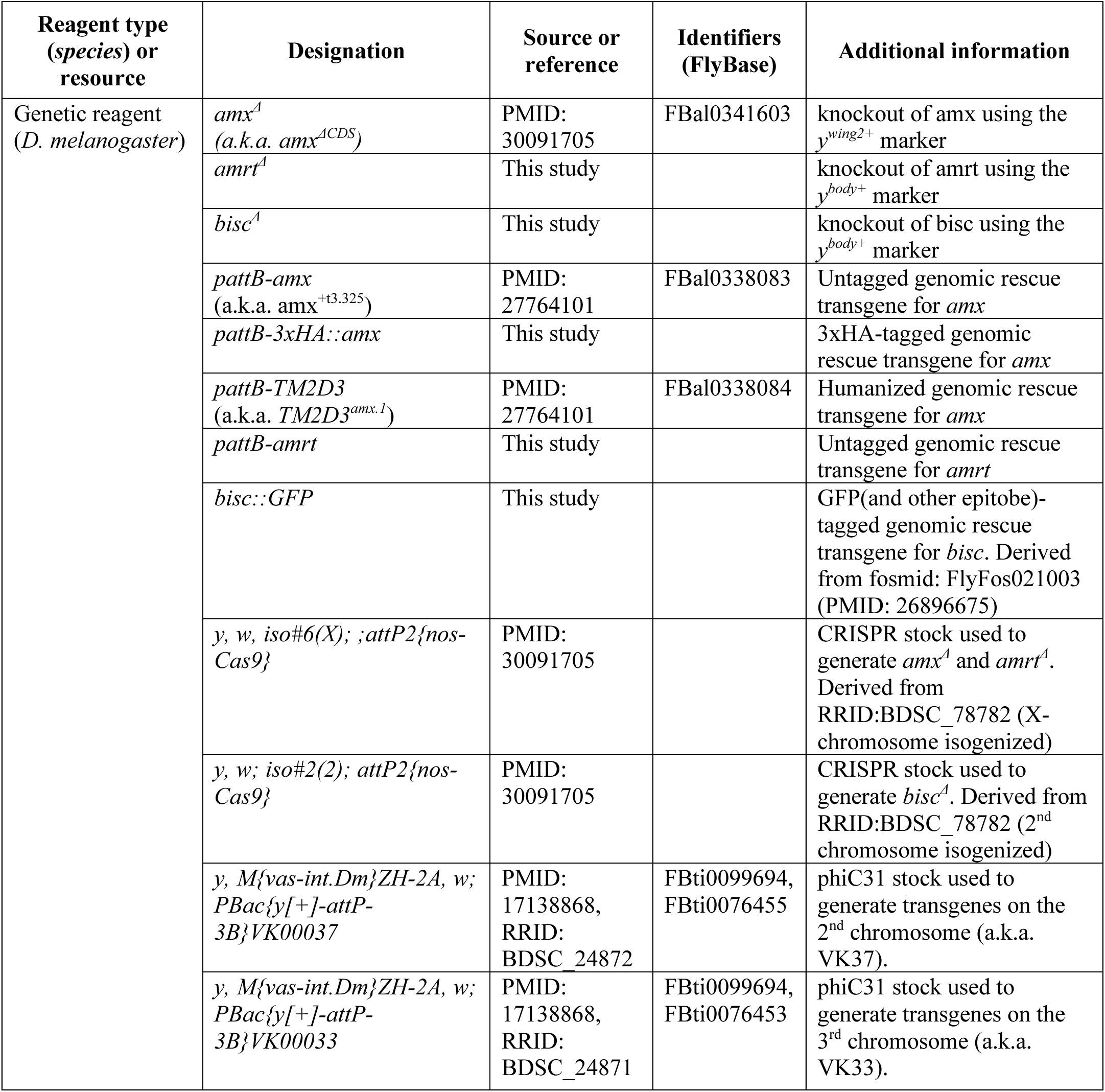
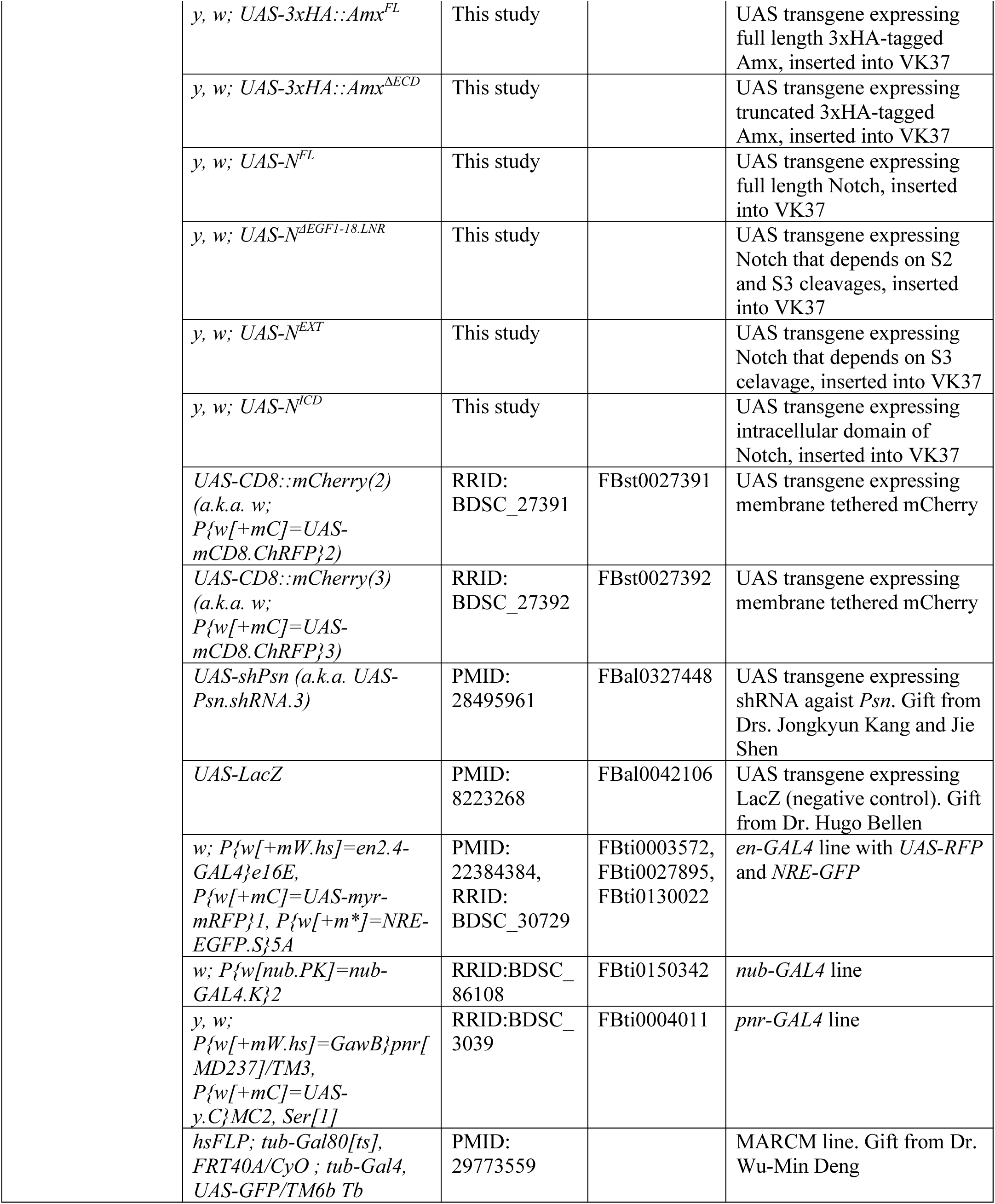
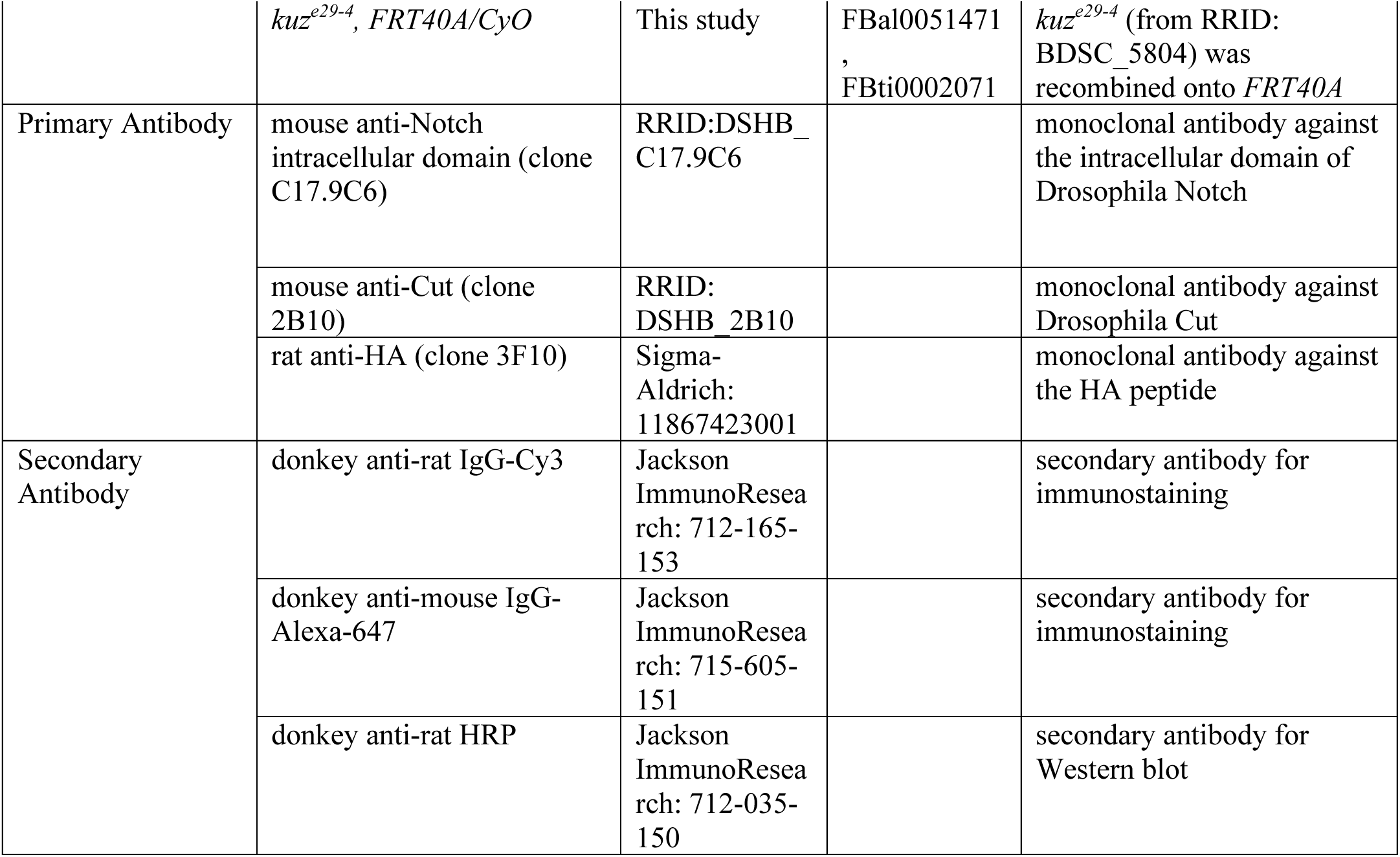

